# ConSeqUMI, an error-free nanopore sequencing pipeline to identify and extract individual nucleic acid molecules from heterogeneous samples

**DOI:** 10.1101/2025.04.03.647077

**Authors:** Adam M. Zahm, Caleb W. Cranney, Alexa N. Gormick, Kathleen E. Rondem, Benjamin Schmitz, Samuel R. Himes, Justin G. English

## Abstract

Nanopore sequencing has revolutionized genetic analysis by offering linkage information across megabase-scale genomes. However, the high intrinsic error rate of nanopore sequencing impedes the analysis of complex heterogeneous samples, such as viruses, bacteria, complex libraries, and edited cell lines. Achieving high accuracy in single-molecule sequence identification would significantly advance the study of diverse genomic populations, where clonal isolation is traditionally employed for complete genomic frequency analysis. Here, we introduce ConSeqUMI, an innovative experimental and analytical pipeline designed to address long-read sequencing error rates using unique molecular indices for precise consensus sequence determination. ConSeqUMI processes nanopore sequencing data without the need for reference sequences, enabling accurate assembly of individual molecular sequences from complex mixtures. We establish robust benchmarking criteria for this platform’s performance and demonstrate its utility across diverse experimental contexts, including mixed plasmid pools, recombinant adeno-associated virus genome integrity, and CRISPR/Cas9-induced genomic alterations. Furthermore, ConSeqUMI enables detailed profiling of human pathogenic infections, as shown by our analysis of SARS-CoV-2 spike protein variants, revealing substantial intra-patient genetic heterogeneity. Lastly, we demonstrate how individual clonal isolates can be extracted directly from sequencing libraries at low cost, allowing for post-sequencing identification and validation of observed variants. Our findings highlight the robustness of ConSeqUMI in processing sequencing data from UMI-labeled molecules, offering a critical tool for advancing genomic research.

**GRAPHICAL ABSTRACT:** 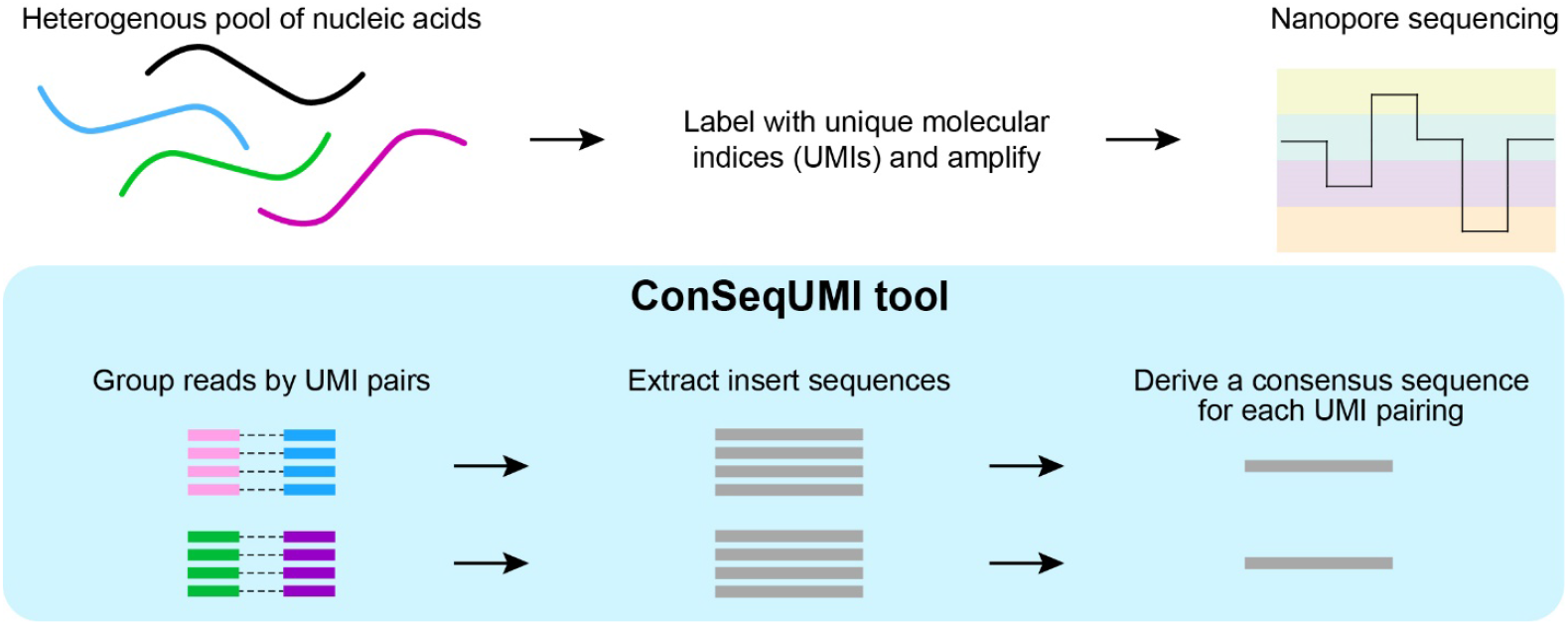

## Introduction

Long-read sequencing technologies utilizing nanopores are highly inaccurate at the single read level relative to short-read methodologies.^1–3^ Nevertheless, the advantages of obtaining full-length sequence information for individual nucleic acids have led to the rapid adoption of these lower-fidelity platforms, especially for the assembly or classification of genomes. Additional applications, such as monitoring viral genome evolution, are well suited for long-read sequencing but the error rates inherent to the platforms limit their implementation to homogenous samples when sequence accuracy is required. Generating accurate consensus sequences using individual reads derived from homogenous samples has been addressed by a number of existing long-read pipelines.^4–13^ Other programs can process long-read data from heterogeneous samples using reference sequences or by making assumptions about sample sequence distributions.^14–18^ A promising conceptual advancement was introduced by Karst *et al*., that employed unique molecular indices (UMIs) to derive consensus sequences from nanopore data without prior knowledge of the sample composition.^19^ This approach allows for the generation of highly accurate sequences by amplifying and sequencing UMI-labeled molecules, facilitating the alignment of multiple reads per molecule. While their method, and those like it, exemplify an excellent starting point for attaining high-accuracy reads, further technical, experimental, and computational development was needed for broader applicability.

Here, we describe a novel long-read sequencing pipeline designed to accurately identify individual, UMI-labeled nucleic acid sequences from mixed pools. Our software is available as a Docker container for easy installation and implementation and can be run from the command line or from a graphical user interface. We have benchmarked our ‘ConSeqUMI’ pipeline using various sample types to confirm the accuracy of generated consensus sequences. We have also developed a cloning plasmid library as an alternative to PCR-based UMI labeling. We then provide a series of experimental vignettes demonstrating the utility of ConSeqUMI across a range of applications including mapping barcoded plasmids, clonal variance in CRISPR-Cas9 editing, and heterogeneity in transgenic and patient derived viral populations. Lastly, we demonstrate how individual clonal isolates can be obtained directly from the sequence library for further analysis or validation.

## MATERIALS AND METHODS

### Reagents

Max Efficiency DH5a Competent Cells (18258012), SuperScript IV One-Step RT-PCR System (12594100), Platinum SuperFi II PCR Master Mix (12369010), Qubit dsDNA High Sensitivity kit (Q33230), DMEM, high glucose (11965092), and Geneticin Selective Antibiotic (10131035) were obtained from ThermoFisher Scientific (Waltham, MA). PrimeSTAR Max DNA polymerase (R045A) was obtained from Takara Bio USA (San Jose, CA). QIAquick PCR Purification Kit (28104), QIAquick Gel Extraction Kit (28506), Plasmid Maxi Kit (12162), QIAshredder (79656), and the AllPrep DNA/RNA Mini Kit (80204) were obtained from Qiagen N.V. (Venlo, The Netherlands). MinION Mk1B Sequencing Device (MIN-101B), Ligation Sequencing Kit (SQK-LSK109) with the Adapter Mix II Expansion (EXP-AMII001), Spot-ON Flow Cell, R9 Version (FLO-MIN106D), MinION Flow Cell, R10 Version (FLO-MIN114), Flongle Flow Cell, R10 Version (FLO-FLG114), Flongle Sequencing Expansion (EXP-FSE002), and the Native Barcoding Kit 96 v14 (SQK-NBD114.96) were obtained from Oxford Nanopore Technologies (Oxford, United Kingdom). dPTP (6H,8H-3,4-Dihydro-pyrimido(4,5-c)(1,2)oxazin-7-one-8-β-D-2’-deoxy-ribofuranoside-5’-triphosphate, Sodium salt) (NU-1119S) was obtained from Jena Bioscience (Jena, Germany). Benzonase (E1014) and OptiPrep Density Gradient Medium (D1556) were obtained from Sigma-Aldrich (Burlington, MA). T4 DNA ligase (M0202) and Gibson Assembly Master Mix (E2611) were obtained from New England Biolabs (Ipswich, MA). LightCycler 480 SYBR Green I Master (04707516001) and KAPA Pure Beads (07983271001) were obtained from Roche (Basel, Switzerland).

### Biological Resources

AAV5_hSyn-eGFP particles and pAAV-hSyn-eGFP plasmid were gifts from Bryan Roth (Addgene viral prep # 50465-AAV5; Addgene plasmid # 50465; http://n2t.net/addgene:50465; RRID:Addgene_50465). The pXR2 (AAV2 Rep/Cap) and pXX6-80 helper plasmids were obtained from the Indiana University School of Medicine National Gene Vector Biorepository (Indianapolis, IN). HEK293T cells (CRL-3216) and HEK-293 cells stably expressing the *Streptococcus pyogenes* Cas9 protein (CRL-1573Cas9) were obtained from American Type Culture Collection (Manassas, VA).

### Statistical Analyses

#### Novel Programs, Software, Algorithms

##### ConSeqUMI: UMI extraction and clustering

In addition to sequencing data files in FASTQ format, ConSeqUMI requires a text file with four lines, representing the four adapters used in the wet lab portion of the protocol. These adapters represent the template-specific and the amplification primer binding portions of the UMI primers or the template-specific sequence and backbone sequences immediately adjacent to the UMI in the pUMI plasmid library. ConSeqUMI utilizes the cutadapt software package to identify these adapters and extract UMI sequences and target sequences from input reads provided in FASTQ format.^20^ The first and last two hundred base pairs of each nanopore read are initially isolated for adapter identification, and are labeled the top and bottom of the read, respectively. The bottom of the read is converted to its reverse complement to reflect the second set of adapters on the complementary strand of the original sequence. Using cutadapt, the front and back linked adapters are used to extract a UMI from the 5’ and 3’ sections of each read. ConSeqUMI recognizes that reverse strands also appear in nanopore data and thus, if cutadapt fails to identify the appropriate linked adapters, the process is repeated on the reverse complement of the entire strand. In this way, the final collection of UMI and target sequences are all representatives of the forward strand of each read. ConSeqUMI implements cutadapt with a 0.2 maximum error rate tolerance for adapter identification.

As an optional setting, ConSeqUMI allows for user-specified UMI lengths to aid in adapter identification. During experimentation, it became clear that some reads are not sequenced to completion, possibly truncating the front adapter of either linked adapter pair. Provided the program knows the expected length of the UMI, an identified adapter internal to the UMI can independently be used to extract the UMI while identifying the start or end of the target sequence. ConSeqUMI allows users to provide the expected UMI length, in which case the front adapters are considered optional, thereby retrieving any prematurely terminated reads containing internal adapters and additional nucleotides equal to or greater than the specified UMI length.

Once adapters are identified and UMIs and target sequences extracted, ConSeqUMI uses the starcode software package to cluster UMIs based on Levenshtein distance similarity.^21^ Starcode offers a significant reduction in time and memory complexity compared to other clustering software for short sequences. The starcode ‘seq-id’ setting attaches lists of original indices to each cluster, retaining memory linking reads to the cluster categorization. UMI pairs are found by running starcode with this setting separately on UMIs extracted from the top and bottom of each read and finding the intersection of indices between their clusters.

##### ConSeqUMI: UMI chimera identification and removal

Once UMI pairs are identified and matched to reads, we follow the same procedure of Karst *et al*. to identify and remove UMI chimeras.^19^ Target sequences are then clustered together for downstream consensus sequence generation. In the final output of the UMI processing step, sequences are saved one FASTQ file per cluster in a bins folder in the output directory. Furthermore, data files regarding read rejection, chimera analysis and starcode output are saved to a data analysis folder in the output directory for further investigation.

##### ConSeqUMI: De novo consensus sequence generation

The pairwise algorithm uses a dual-method approach to generating consensus sequences from bins of target sequences. In the first stage of the pairwise method, a preliminary consensus sequence is generated using a custom frameshift algorithm. After buffering the ends of the binned sequences with spaces to match the maximum (central vector) length, the algorithm extracts a ten base pair window from the front of each sequence. The most common sequence is established as the front end of the consensus sequence. The algorithm determines the most common eleventh base pair for all sequences that contain the initial ten base pairs. Each base pair is then added sequentially, searching for the base pair immediately following the last ten identified base pairs of the consensus sequences. These ten base pairs are searched within a twenty base pair window that shifts for every identified base pair. This method has several flaws in determining the final consensus sequence, such as possibly skipping or duplicating base pairs in homopolymeric regions, but generates a respectable reference sequence. This method is also used for providing a reference to the medaka consensus algorithm.

In the second stage of the pairwise method, the preliminary consensus sequence generated in the first stage is adjusted to more accurately reflect binned sequence commonalities. A pairwise alignment between the preliminary consensus sequence and each binned sequence is performed, where individual discrepancies are saved and collected. The most common discrepancy between all binned sequences is then applied to the consensus sequence. This process is repeated so long as each added discrepancy increases the average alignment score. When an added discrepancy decreases the average, the discrepancy is ignored and the prior consensus sequence is considered the true consensus sequence.

Provided a directory of FASTQ files wherein each file contains reads expected to coincide with a single consensus, such as the bin directory output of the UMI processing step, ConSeqUMI generates a single FASTA file containing the consensus sequence for each FASTQ file. By default, ConSeqUMI only generates consensus sequences for directories/clusters with 50 or more reads. However, this benchmark can be adjusted by the end user, as the sequencing error rate can differ significantly between sequencing protocols and runs. To help researchers establish a benchmark level appropriate for their own data, ConSeqUMI includes a bootstrapped analysis workflow in addition to the UMI processing and consensus generation workflows. This setting extracts a set of binned reads, such as an output FASTQ file of the UMI processing step, and iteratively and randomly extracts subclusters incrementing at a user-defined size. A consensus sequence is generated from each subcluster, and subsampling at each size occurs a user-defined number of times. While both the subsampling step size and iterations of each step size can be set by the end user, the default values are 10 and 100, respectively. The Levenshtein distance between consensus sequences generated for each subcluster and the one generated from the full cluster are then collected and returned in a comma-delimited file for benchmark cutoff analysis. Users can then set read number cut-off in ConSeqUMI suitable from the given experiment to achieve the desired accuracy.

### Web Sites/Data Base Referencing

The ConSeqUMI package is available on GitHub (https://github.com/JGEnglishLab). MinKNOW software version 23.11.7, Guppy, and Medaka (https://github.com/nanoporetech/medaka) were obtained from Oxford Nanopore Technologies (Oxford, United Kingdom). The following tools were accessed using Galaxy: Filter BAM, Collapse and Trim sequences, and NextClade.^22–24^ We also utilized the following software: Minimap2, Tablet, SAMtools, Qualimap, Clustal W, Clustal Omega, Cutadapt, and SANKEY_PLOT Stata module.^20,25–32^

### Primer-based UMI labeling

One important consideration for primer-based labeling is the input sample amount. When too many molecules are UMI-labeled and amplified, the sequencing read depth is insufficient to generate accurate consensus sequences. We have found that approximately 10^5^-10^6^ input molecules is a good target amount per sample, but the ideal number will depend on the sequencing format (MinION, GridION, etc.) and the number of samples being pooled for sequencing. We have had success controlling input molecule number at the time of UMI labeling (i.e. input amount in the UMI PCR) or at the time of labeled-product amplification by diluting the UMI PCR output. What follows is a general description of the procedures we used to label and amplify target molecules using PCR. Methodologies utilized in specific experiments are described in relevant sections below. Primer sequences are listed in Table S1. PCR reagents and conditions used are listed in Table S2.

UMI-labeling was performed using either Platinum SuperFi II PCR Master Mix or PrimeSTAR Max DNA polymerase for two cycles of PCR according to manufacturer’s instructions. Reactions were then purified to remove UMI primers using either KAPA Pure Beads at 0.6X or the QIAquick PCR Purification Kit, both according to manufacturer’s instructions. Purified products were then amplified with PCR using primers binding externally to the UMI labels. Following the amplification PCR, products were examined via agarose gel electrophoresis. Purification of products was then performed by gel extraction or by column purification of the remaining reaction volume using the QIAquick Gel Extraction Kit or QIAquick PCR Purification Kit, respectively.

### Nanopore library preparation and sequencing

For the initial barcoded amplicon experiment, the UMI-labeled amplicons were prepared for nanopore sequencing using the Ligation Sequencing Kit with the Adapter Mix II Expansion according to protocol NBA_9093_v109_revJ_12Nov2019 using KAPA Pure Beads in place of AMPure XP beads. Sequencing was performed with the Spot-ON Flow Cell, R9 version, using the MinION sequencer. Raw FAST5 trace files were basecalled and demultiplexed using the Guppy software package with ‘sup’ accuracy and without adaptive sampling. For all other experiments, UMI-labeled amplicons or amplicons from pUMI were prepared for nanopore sequencing using the Native Barcoding Kit 24 v14 (SQK-NBD114.24) according to protocol NBA_9168_v114_revE_15Sep2022, using KAPA Pure Beads in place of AMPure XP beads. Sequencing was performed with MinION Flow Cells (R10 version) or Flongle Flow Cells (R10 version) using the MinION sequencer with live read basecalling by the MinKNOW software using ‘sup’ accuracy and without adaptive sampling. We used the default MinKNOW read quality thresholds to classify failed and passed reads. All failed reads were omitted from processing, while all passed reads were included.

### Nanopore sequence data processing

All processing of sequencing data was performed using the ConSeqUMI package. UMI processing was typically performed using the optional ‘UMI length’ flag set to 18. Medaka models r941_min_sup_g507_model and r1041_e82_260bps_sup_g632_model were downloaded from the medaka Github. Resulting consensus sequence outputs in FASTA format were analyzed as applicable. Consensus sequences were aligned to reference sequences using Minimap2 and resulting .SAM files were visualized using Tablet, while .BAM and .pileup files were generated with SAMtools. Commands and flags used are listed in Table S5. Quantification of indels was performed using Qualimap and quantification of mismatches was performed using a previously published script.^33,34^

### Generation and sequencing of amplicons containing unique internal barcodes

UMI-containing PCR primers were used to generate 1440 bp amplicons from a pooled plasmid library containing approximately 1.7×10^5^ unique 24 nucleotide barcodes (minCMV fraction; ‘barcode library’).^35^

### Plasmid pool amplicon generation and sequencing

Fourteen pcDNA3.1-derived plasmids with unique expression cassettes were pooled (‘pooled plasmid library’; sequences listed in Table S3). UMI-containing PCR primers targeting universal backbone sequences were used to generate amplicons of varying length. Approximately 10^5^ or 10^6^ plasmids were UMI-labeled independently and in triplicate by two cycles of PCR using Platinum SuperFi II PCR Master Mix. Reactions were purified with magnetic beads and eluted in 10 uL of water. Amplicons were then amplified with 35 cycles of PCR using Platinum SuperFi II PCR Master Mix. Products were isolated by column purification. Primer sequences and reaction parameters are listed in Tables S1 and S2, respectively. The sequences of the plasmids in the pool, as well as the PCR target regions, are listed in Table S3.

To assess reproducibility of UMI labeling, the consensus sequences for all clusters regardless of read depth were derived for each technical replicate using ConSeqUMI with the pairwise method and mapped to the pooled plasmid library sequences. Reads mapping to each plasmid of the pool were tallied using Tablet. Benchmarking was performed for each consensus sequence generation algorithm with a sub-cluster sample interval size of one read with 50 iterations at each interval up to size 30. For quantification of mismatches and indels, we limited analyses to pairwise method consensus sequences for clusters with a minimum of 17 reads. To tabulate reads with and without errors, BAM files derived from consensus sequences produced by the pairwise method were filtered using the Filter BAM tool on Galaxy with a tag value of ‘NM:>0”.

### Comparing read accuracy of flow-cell and reagent versions

To compare read accuracy for the R9 and R10 version MinION flow cells, we aligned individual reads of a single large cluster from both the barcode amplicon and plasmid pool data sets to the respective consensus sequences derived for those clusters using the pairwise method. Resulting pileup and BAM files were used to count mismatches and indels as described above.

### Long-term bacterial outgrowth mutagenesis

A glycerol stock of a clonal bacterial culture harboring our pcDNA3.1-mGreenLantern plasmid was used to inoculate two mL of lysogeny broth (LB). This inoculated LB was cultured for 48 hours at 37°C with shaking. One microliter of this culture was then used to inoculate a fresh two mL of LB, which was then cultured for 48 hours at 37°C with shaking. After the second 48-hour period, the culture was split to both establish a new glycerol stock and to isolate plasmid DNA using the QIAprep Spin Miniprep Kit (Qiagen). This process was repeated for a total of six rounds (‘weeks’). Primer-based UMI labeling of plasmids was performed with approximately 10^6^ input molecules using the PCR conditions outlined in Table S2. Initial PCR products were purified using the QIAquick PCR Purification Kit and eluted in 25 uL of EB buffer. The elution was then diluted 1:5 and 1 uL of this dilution was used as input in the 20 uL amplification PCR.

Consensus sequences were derived using the pairwise method for all clusters with 25 or more reads and then aligned to the input plasmid sequence using minimap2. Quantification of mismatches and indels was performed as described above. Correlation between bacterial culture time and observed error rates was assessed using Spearman’s rank correlation test. Pearson’s chi-square was used to test differences in read depth between clusters with and without errors.

### Amplicon mutagenesis with repeated PCR and with a nucleoside triphosphate analog

To introduce random mutations in a DNA template, we first amplified a 3296 bp fragment from pcDNA3.1 containing a human GNA15 CDS using PrimeSTAR Max DNA polymerase or Platinum SuperFi II Green PCR Master Mix according to manufacturer’s instructions. Products were purified using the QIAquick PCR Purification Kit and subjected to additional rounds of PCR and purification. Each reaction was performed using 0.5 ng of template DNA. Products from each round of PCR, as well as the original plasmid template, were UMI-labeled with two cycles of PCR using Platinum SuperFi II Green PCR Master Mix. Amplification of UMI-labeled molecules was performed with the respective polymerase reagents used in the repeated PCRs. Consensus sequences were generated with the pairwise method for all PrimeSTAR-derived clusters with 20 or more reads and all SuperFi-derived clusters with 10 or more reads. Mismatches and indels were quantified as described above. Estimations of PCR error rates were determined using linear regression. Tests of correlation between mutation rates and PCR cycles were performed with cluster numbers as probability weights.

To introduce random mutations in a DNA template using a nucleoside triphosphate analog, we amplified a 3296 bp fragment from pcDNA3.1 containing a human GNA15 CDS (pcDNA3.1_GNA15) using PrimeSTAR Max DNA polymerase in the presence of varying concentrations of dPTP. Reactions were performed according to manufacturer’s instructions. Products were purified using the QIAquick PCR Purification Kit and quantified using the Qubit dsDNA HS kit. Products from each reaction were UMI-labeled with two cycles of PCR using Platinum SuperFi II Green PCR Master Mix. Amplification of UMI-labeled molecules was performed using PrimeSTAR Max DNA polymerase. Consensus sequences were generated with the pairwise method for all clusters with 10 or more reads. Mismatches and indels were quantified as described above. Reference, consensus, and Sanger-derived sequences were aligned using Clustal W.

### AAV generation and sequencing

An AAV-ITR plasmid expressing eGFP under control of a CMV or EF1α promoter, a U6 promoter and an empty sgRNA cassette was synthesized by Genscript. A DNA fragment containing a U6 promoter and an IRF3-targeting sgRNA cassette was synthesized as a gene block from Integrated DNA Technologies (IDT) and was inserted upstream to the eGFP of the AAV-ITR plasmid using Gibson Assembly (NEB, E2611S) to create the pITR-IRF3_sgRNA plasmid. Guide RNA sequences of interest were selected from the Brunello Library (Addgene, #73179). AAV2 virions were prepared using helper-free methods described in Grieger *et al*.^36^ Briefly, HEK293T cells were seeded into 15-cm dishes at a density of 7×10^6^ cells per dish and maintained at 5% CO_2_ in Dulbecco’s modified Eagle medium supplemented with 10% fetal bovine serum. When the cell density reached 70%, calcium phosphate transfections were performed by adding pXR2 (Rep/Cap), pITR-sgRNA, and pXX6-80 helper plasmids in a 1:1:3 ratio.

AAV2 virions were harvested from cellular extracts 72 hours after transfection using standard methods. Briefly, cells were scraped and collected into a 50 mL conical and spun at 500 x g to remove cellular debris. Pelleted cells were resuspended in a hypotonic buffer, and incubated on ice for ten minutes. Cells were then lysed using a combination of dounce homogenization and sonication (Branson Sonifier 250) for one minute (50% duty cycle, output setting 3). Cell lysates were then clarified by centrifugation at 2000 x g for 10 minutes. AAV virions were isolated by loading the clarified supernatant on to a manually stacked iodixanol density gradient and spun overnight at 100,000 x g. Genome-packaged AAV fractions were identified by RT-qPCR, using probes targeting the packaged ITR-sgRNA genomes, pooled, and concentrated using an Amicon Ultra filter (50K MWCO) to remove residual iodixanol. Purified AAVs were then flash frozen in DPBS and stored at -80°C until the time of assaying.

Genomic DNA was isolated from AAV virions using phenol/chloroform according to standard procedures and DNA pellets were resuspended in TE buffer. Prior to UMI-labeling of genomic DNA from our AAVs preparations, we performed an initial round of PCR with 30 cycles and column purified the reactions. Isolated DNA from the commercial AAV5 preparation UMI-labeled without preamplification using approximately 10^5^ input molecules. Primers and PCR conditions are detailed in Tables S1 and S2, respectively. Consensus sequences for the commercial AAV5 sample were derived using the pairwise method for clusters with 24 or more reads. For the generated AAV2 virions with the CMV or EF1 expression cassette, consensus sequences were derived for clusters with a minimum of 12 or 14 reads, respectively.

### UMI plasmid generation

To generate the pUMI library, two cycles of PCR using primers with degenerate, patterned UMI regions were used to introduce tandem UMIs separated by a EcoRI recognition site at the MCS of pUC19. After column purification to remove primers, products were amplified via PCR using introduced primer binding sites external to the UMI regions. Resulting amplicons were digested with EcoRI and ligated in a 200 µL reaction using T4 DNA ligase. The ligation mixture was then transformed into 400 uL of Max Efficiency DH5a Competent Cells. Dilutions of the recovery media were plated to estimate total diversity and the remaining volume was cultured in 2X YT media overnight at 30°C. Plasmid DNA was then isolated using the Plasmid Maxi Kit.

### CRISPR/Cas9-induced genomic mutations

HEK-293 cells stably expressing the *Streptococcus pyogenes* Cas9 endonuclease (SpCas9) were seeded into 6-cm dishes at a density of 2×10^5^ cells per dish in Dulbecco’s modified Eagle medium supplemented with 10% fetal bovine serum and 200 ug/mL Geneticin. Titers of purified AAV2 virions harboring a U6-IRF3-sgRNA cassette were determined via qPCR using LightCycler 480 SYBR Green I Master. At the time of seeding, virions were added on top of cells (10^3^ genomes per cell) and dishes were cultured overnight in a 37°C cell culture incubator. The following day, the media was exchanged and the cells were maintained for a period of seven days. Cells were then scraped from the dishes and pelleted by centrifugation. Genomic DNA was isolated from cell pellets using QIAshredders and the AllPrep DNA/RNA Mini Kit. Target regions of genomic DNA were amplified using the Platinum SuperFi II Green PCR Master Mix to introduce EcoRI restriction sites (for compatibility with pUMI. Products were digested and ligated to the pUMI backbone. Following transformation and culture of chemically competent DH5a cells, colonies were scraped from agar plates and plasmids were isolated using the QIAprep Spin Miniprep Kit. Fragments containing the inserts and flanking UMIs were liberated from plasmids using PCR with primers binding to the M13 sites flanking the inserts. Consensus sequences were generated for all clusters with six or more reads using lamassemble and mapped to the human IRF3 target region using minimap2. Mapped reads were trimmed to 100 nucleotides centered on the predicted SpCas9 cleavage site using Trim sequences of the FASTX-Toolkit in Galaxy. Resulting trimmed reads were collapsed using Collapse in Galaxy.

### SARS-CoV-2 spike glycoprotein CDS processing

SARS-CoV-2 genomic RNA samples isolated from saliva were obtained from Dr. David O’Connor, University of Wisconsin-Madison.^37^ A 6162 bp region containing the full spike glycoprotein CDS was reverse transcribed and amplified using the SuperScript IV One-Step RT-PCR System according to manufacturer’s instructions. Primers were designed to target regions that were not mutated in Delta and Omicron variants. Amplicons were isolated via agarose gel electrophoresis and quantified with the Qubit dsDNA HS Assay Kit. UMIs were added to undiluted amplicons with two cycles of PCR using the Platinum SuperFi II Green PCR Master Mix. Reactions were purified using the QIAquick PCR Purification Kit and eluted in 20 uL EB. Products were diluted, amplified with PrimeSTAR Max DNA Polymerase, and isolated via agarose gel electrophoresis. Primer sequences and reaction parameters are listed in Tables S1 and S2, respectively.

Cluster consensus sequences generated by ConSeqUMI were trimmed using Cutadapt to produce spike glycoprotein CDS sequences. Resulting sequences were collapsed by sequence identity using Collapse and were then classified using NextClade. Uncollapsed reads were aligned with Clustal Omega and parsed for haplotype/linkage disequilibrium analyses. Translations of Spike CDS sequences were aligned and clustered (neighbor-joining) using Clustal Omega. Sankey plots were generated using the SANKEY_PLOT Stata module.

### Target retrieval from library pools

PCR primers corresponding to the UMI sequences of individual clusters of interest were used to isolate transgenes from the heterogeneous amplicon pools. PCR was performed using the Platinum SuperFi II Green PCR Master Mix according to manufacturer’s instructions. Amplicons were isolated by agarose gel electrophoresis and extracted using the QIAquick Gel Extraction Kit. Regions of interest were sequenced via Sanger sequencing. Primer sequences and PCR conditions are listed in Tables S1 and S2, respectively.

## RESULTS

### ConSeqUMI software

The high error rates of nanopore-based sequencing technologies can be overcome with sufficient read depth such that errors can be polished from individual reads to derive an accurate consensus sequence. However, read polishing is not possible when the sample being sequenced is heterogeneous in identity. Karst *et al*. proposed an elegant solution to this problem: uniquely label molecules to be sequenced with dual unique molecular indices (UMIs) and then amplify these labeled molecules prior to sequencing.^19^ In this manner, multiple sequenced reads can be obtained for individual molecules present in heterogeneous samples. We have optimized this approach and developed a software package, ConSeqUMI, to cluster UMI-labeled amplicons and generate consensus sequences from as few as 10 reads per molecule. Initially, long-read sequencing data files provided in FASTQ format are parsed for adapter sequences flanking UMI regions. Reads with matching UMI pairs are then binned together and a consensus sequence is derived for each individual cluster using the clustered reads (Fig. 1). ConSeqUMI offers three methods for consensus sequence generation: the existing consensus generation software packages medaka and lamassemble, as well as a novel technique we introduce, the “pairwise” method (see Methods).^38^ Notably, the output bins produced by ConSeqUMI are amenable to consensus sequence generation using alternative tools, if desired. To help identify read depth thresholds necessary for accurate consensus sequence generation, we developed a benchmarking script in ConSeqUMI that subsamples reads from a single cluster in a sequencing run, iteratively generates consensus sequences from sequentially larger subsampling bins, and compares these to the consensus sequence derived using all available reads of the cluster. This benchmarking tool can predict the minimum required cluster size for perfect read accuracy for any sequencing run, allowing the user to maximize their sequenced genomes. The ConSeqUMI package is available on GitHub (https://github.com/JGEnglishLab).

**Figure 1.**
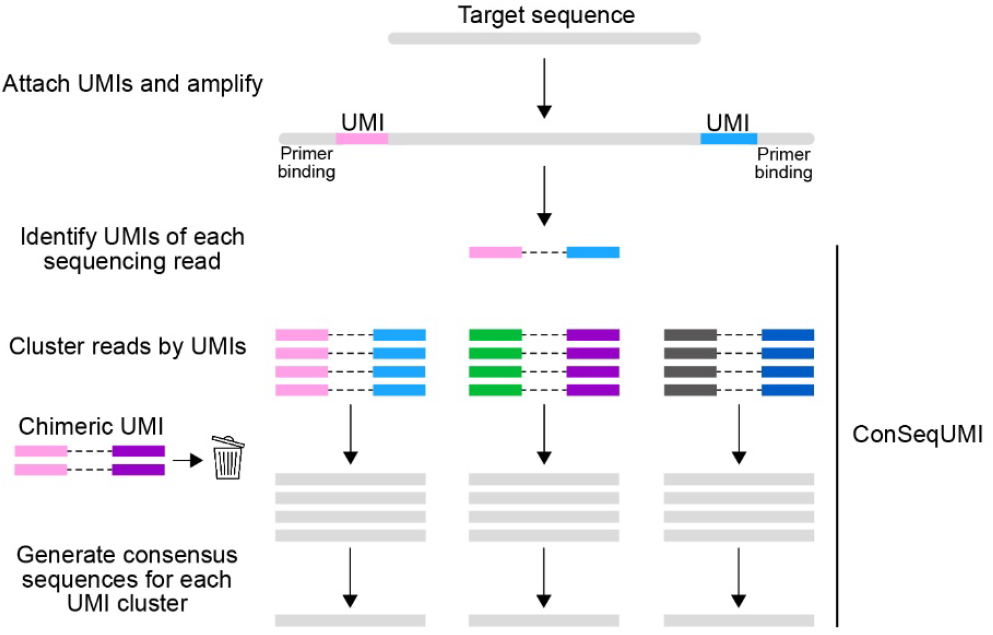
Overview of target molecule amplification and sequencing data processing by ConSeqUMI.

### Chimeric read identification

One potential complication of primer-based UMI labeling is the generation of chimeric amplicons, in which an original UMI-tagged molecule has one or both UMIs replaced by residual UMI primers carried over into the amplification PCR.^19^ ConSeqUMI automatically identifies and discards clusters containing chimeric UMIs, in which one or both of a cluster’s UMIs are also found in other clusters with higher read depth. There remains a possibility in which an individual molecule gets both original UMIs replaced with two new UMIs during amplification and that these new UMIs do not appear in other clusters. In this case, the relative abundance of the molecule within the target pool would be artificially increased. To determine if these ‘double chimera’ events occur, we generated UMI-labeled amplicons from a plasmid pool containing approximately 1.7×10^5^ unique 24-nt barcodes (Fig. 2A).^35^ If a single barcode was found to be flanked by more than two pairs of unique UMIs, this would likely reflect a double chimera labeling due to the complexity of the template library (Fig. S1). We used a MinION R9 flow cell and associated reagents to sequence UMI products derived from approximately 10^5^ or 10^6^ input molecules to identify a target number of molecules that will maximize the number of clusters with read depths necessary for accurate sequence identification.

**Figure 2.**
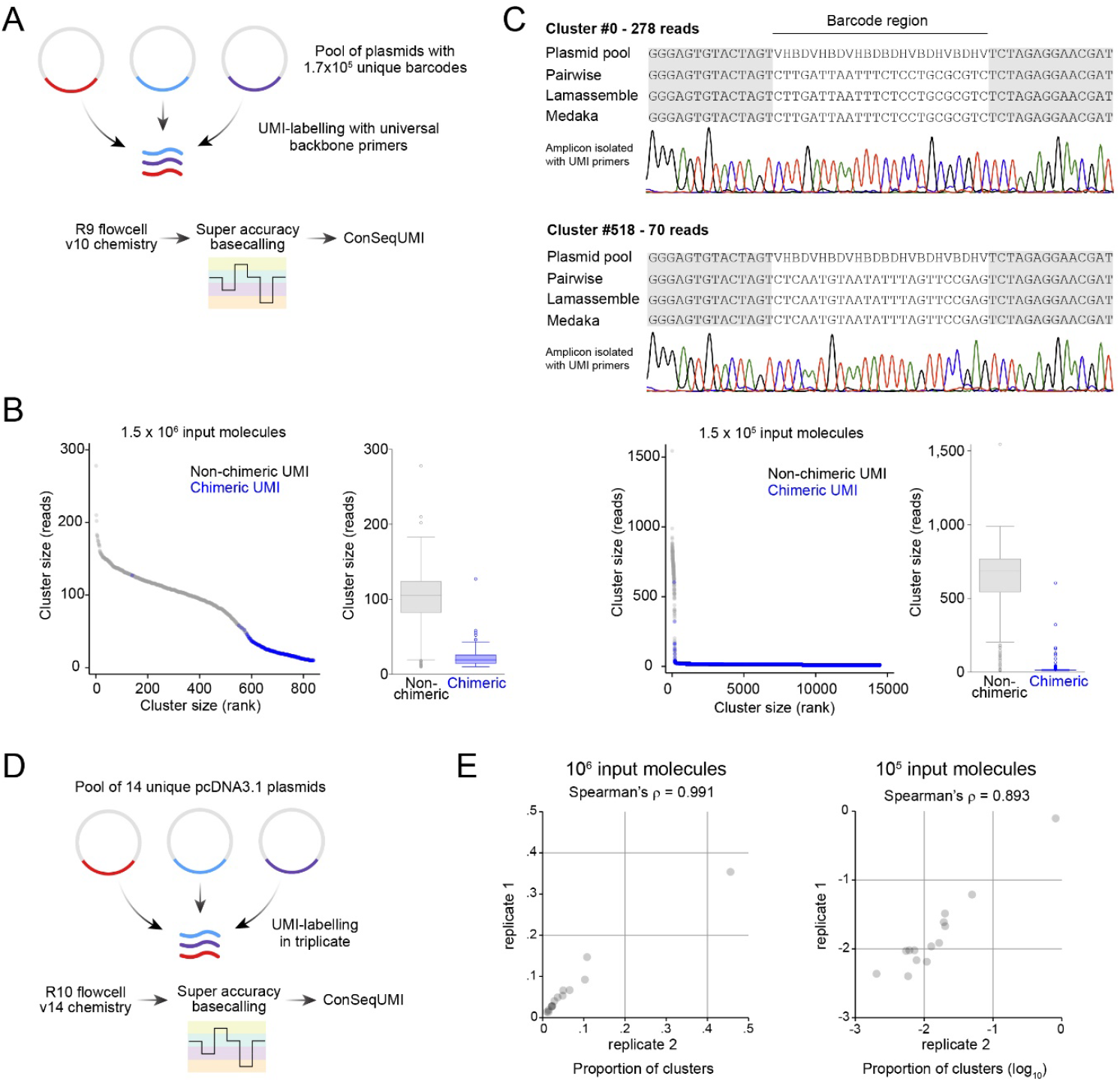
Benchmarking ConSeqUMI using barcoded amplicons and a pool of heterogeneous plasmids. (**A**) Chimeric read experiment schematic. UMI tags were appended to flanking regions of the degenerate barcode of a pool barcoded plasmids using two cycles of PCR with universal primer binding sites. Following bead cleanup, amplicons were further amplified with additional PCR cycles using appended constant regions as primer binding sites. Amplicons were prepared for nanopore sequencing using v10 chemistry native barcoding and sequenced on a MinION sequencer using an R9 flow cell. **B**) Cluster sizes in reads of non-chimeric and chimeric UMI pairings for 1.5 × 10^6^ (top panel) or 1.5 × 10^5^ (bottom panel) input molecules following ConSeqUMI processing. Clusters containing chimeric UMIs are displayed in blue. Inset box plots summarize the data of the quantile plots. **C**) Isolation of two transgenes from heterogeneous pools via PCR using UMIs as primer binding sites. Abridged cluster consensus sequences for each ConSeqUMI algorithm are shown below the template sequence region containing degenerate nucleotides, with the constant backbone regions in gray. Chromatograms display the results of Sanger sequencing of the resulting amplicons. **D**) Plasmid pool experiment schematic. UMI tags were appended to variable regions of a pool of 14 pcDNA3.1-based plasmids using two cycles of PCR with universal primer binding sites on the pcDNA3.1 backbone. Following bead cleanup, amplicons were further amplified with additional PCR cycles using appended constant regions as primer binding sites. Amplicons were prepared for sequencing using v14 chemistry native barcoding and sequenced on a MinION sequencer using an R10 flow cell. **E**) Replicate UMI-labeled plasmid pool samples were sequenced and clustered using ConSeqUMI. Primer-based UMI addition showed consistent cluster identity proportions across replicates per input amount.

After identifying unique UMI pairs, binning reads into clusters, and extracting the amplicon barcode sequences, we calculated the prevalence of chimeric UMIs. In the sample with 10^6^ input molecules, ConSeqUMI identified 574 clusters with 50 or more reads (Fig. 2B). Of these, four (0.7%) were discarded because of a chimeric UMI pairing. In the remaining 570 clusters, we identified two 24-nt inserts that were each present in two separate clusters. In the sample with 10^5^ input molecules, ConSeqUMI identified 193 clusters with 50 or more reads (Fig. 2B). Of these, eight (4.1%) were discarded because of a chimeric UMI pairing. From these results, we conclude that PCR-based UMI labeling produces a low level of chimeric amplicons and that double chimeric events are extremely rare and are unlikely to affect conclusions regarding relative molecule abundance.

Next, we utilized the ConSeqUMI benchmarking script to evaluate the accuracy of each of the three consensus generation algorithms using the two clusters with the highest read depth. We generated subclusters of all sizes, from one read to the maximum available reads, by randomly sampling from all available reads, generated consensus sequences for each of these subclusters, and then calculated the Levenshtein distances of the consensus sequences to that of the full cluster. We performed fifty iterations at each subcluster size for each of the consensus generation algorithms. For both sampled clusters, the Lamassemble algorithm outperformed the Medaka and Pairwise algorithms, as it generated consensus sequences that were nearest to the full cluster consensus sequences (Fig. S2A) and had the greatest number of iterations showing a Levenshtein distance of zero (Fig. S2B). Because the Levenshtein distances continued to decrease with increased subcluster size across much of the range, we concluded that some of our clusters did not contain the read depth necessary to generate a perfect consensus sequence across the entire amplicon.

Because each input DNA strand is labeled with two unique UMIs, we reasoned that specific target sequences could be extracted from heterogeneous amplicon pools using PCR primers of sequence corresponding to the targets’ flanking UMIs. To test this hypothesis, we first performed PCR to selectively amplify two separate target sequences identified by ConSeqUMI in our barcoded template experiment and then Sanger sequenced the resulting amplicons. Notably, the consensus barcode sequences derived by each of the ConSeqUMI algorithms for the selected targets were in agreement (Fig. 2C). The barcode regions of the amplicons from the UMI-specific PCRs validated the consensus sequence predictions of ConSeqUMI and confirmed that specific individual target sequences can be isolated from heterogeneous pools of UMI-labeled molecules.

### Reproducible reference-free sequencing of heterogeneous samples

New analysis pipelines can process Nanopore sequencing data containing heterogeneous inputs without the need for barcoding and demultiplexing.^18,39^ However, these approaches require reference sequences for the alignment and identification of reads or derive ‘haplotype’ sequences under the assumption that reads of similar sequence are derived from identical molecules. The ConSeqUMI pipeline does not require users to supply reference sequences and generates consensus sequences of individual molecules independently. Thus, in theory, it can identify fully divergent nucleotide sequences present within a single input sample. To test this hypothesis, and to determine whether UMI labeling using degenerate primers consistently labels target molecules proportionally, we unevenly pooled 14 plasmids with unique inserts (range of insert size: 404-1884 bp) in a common backbone (Table S3). We labeled approximately 10^5^ or 10^6^ pooled molecules independently and in triplicate, nanopore sequenced the amplicons using an updated flow cell (R10) and reagents, and mapped the resulting cluster consensus sequences from ConSeqUMI to the input ‘genome’ i.e. the sequences of plasmid inserts (Fig. 2D). The proportion of cluster consensus sequences mapping to the 14 unique inserts (regardless of cluster read depth) were strongly correlated across replicates, demonstrating the reproducibility of primer-based UMI labeling combined with ConSeqUMI processing (Fig. 2E, S3A). Of note, the correlation coefficients between replicates of 10^5^ input molecules were lower, suggesting that the precision of target proportionality identified by primer-based labeling is dependent on the number of input molecules. Thus, in scenarios where proportionality is important, we suggest attempts be made to label the largest number of input molecules that will provide adequate cluster read depth.

The upgraded flow cell and reagent kit used in this experiment greatly reduced the number of sequencing reads necessary to derive accurate consensus sequences. Sub-sampling sequence reads from two separate clusters and performing 50 iterations of consensus generation at sample sizes from 1-30 reads found that lamassemble generated the correct consensus sequence for 100% of iterations using 21 or more reads (Fig. S3B,C). Our pairwise method identified the correct consensus sequence for all iterations using 14 or more reads. Meanwhile, medaka was unable to consistently generate the correct consensus with 30 sub-sampled reads. To determine whether the reduced cluster density requirement observed with the updated flow cell and reagents was due to lower basecall error rates, we mapped the individual reads of single clusters from this experiment, and from the barcoded amplicon experiment using the R9 flow cell, to the respective cluster consensus sequences and quantified mismatches, insertions and deletions. As expected, the R10 flow cell and associated reagents showed lower rates for all mismatch types as well as fewer indels per read (Fig. S3D). Basecall errors were more likely to resemble transition rather than transversion mutations, suggesting the Oxford Nanopore platform has greater difficulty distinguishing between two purines and between two pyrimidines, as observed previously.^40^

Next, we sought to verify that consensus sequences generated by ConSeqUMI matched the expected plasmid templates. We generated a consensus sequence for each cluster with 17 or more reads using the pairwise method, aligned these sequences to the expected sequence of the plasmid templates, and tabulated mismatches, insertions, and deletions. Of 15,782 total cluster sequences across the six samples, 82 (0.52%) of consensus sequences contained at least one error (in total: 74 mismatches, seven insertions, and 5 deletions). We noted that mismatches were more often found at reference sequence guanines or cytosines (Fisher’s exact, P<.0001) (Fig. S4A). These observed mismatch types suggest that these consensus sequence errors are not a result of Nanopore basecall errors, as they did not mirror the pattern of mismatch types observed in our analysis of individual reads (Fig. S4B). We hypothesized these errors: (a) resulted from ConSeqUMI consensus generation for clusters with low read depth, (b) represent mutations acquired during clonal outgrowth of plasmids, or (c) were PCR-induced mutations during UMI labeling.

If the observed errors were a result of insufficient read depth, we would expect the clusters with errors to contain fewer reads than clusters without errors. Thus, we tabulated the read depth for all clusters with and without errors and compared these distributions (Fig. S4C). In one of the six samples, clusters with errors had significantly fewer reads than error-free clusters, whereas five samples showed no difference between groups. Nevertheless, in each sample, correct consensus sequences were derived for clusters that contained as few or fewer reads than those clusters whose sequences contained errors. As a group, there was no significant difference in read depth between clusters with or without errors (Wilcoxon rank sum z=.0434, P=0.9654). Furthermore, the proportion of sequences with errors was variable across plasmid templates, ranging from 0-3.9% (Fig. S4D). Thus, we concluded that read depth was not a major contributing source of error, in agreement with the read depth benchmarking we performed using ConSeqUMI.

### Clonal plasmid stocks do not acquire mutations during outgrowth

To determine whether plasmids acquire mutations during bacterial cell outgrowth at a rate that would explain mismatch errors observed in our plasmid pool experiment, we cultured clonally-derived *Stbl3* bacterial cells harboring one of the plasmids from our mixed pool experiment for six weeks in liquid media and assessed plasmid mutations across longitudinal samples (Fig. 3A). We generated UMI-labeled amplicons that included the plasmid’s origin of replication, antibiotic resistance cassette, and a fluorescent protein expression cassette. We hypothesized that mutations accumulating during bacterial cell proliferation would occur at lower rates within the origin of replication and antibiotic resistance regions compared to the fluorescent protein cassette. Mismatch errors followed a similar pattern as we observed in the plasmid pool experiment - occurring more often at reference sequence guanines or cytosines (Fisher’s exact, P<.0001) (Fig. 3B). However, we did not observe an increase in mutation frequency with increased bacterial culture time (Spearman’s ρ=-.0904, P=.8279), suggesting these mutations were either present prior to the experiment or they were introduced by PCR following plasmid purification, or both. All mismatches and indels occurred at unique positions and there were no differences in mutation rates within or outside of the origin of replication and ampicillin resistance gene regions (Pearson’s chi square=.0162, P=.8989). (Fig. 3C). Furthermore, three of the four mismatches detected in the ampicillin resistance gene are missense mutations. As in our plasmid pool experiment, there was no difference in cluster size between clusters with errors and those without (Fig. S5). The percentage of clusters across all time points containing a mismatch was similar to the percentage observed for this plasmid in the plasmid mix experiment (0.91% and 0.35%, respectively; Fisher’s exact test P=0.505). These results demonstrate that plasmids do not acquire mutations during bacterial outgrowth at a frequency detectable at this sequencing depth, aligning with historic measurements of bacterial mismatch repair and mutagenesis frequency.^41,42^

**Figure 3.**
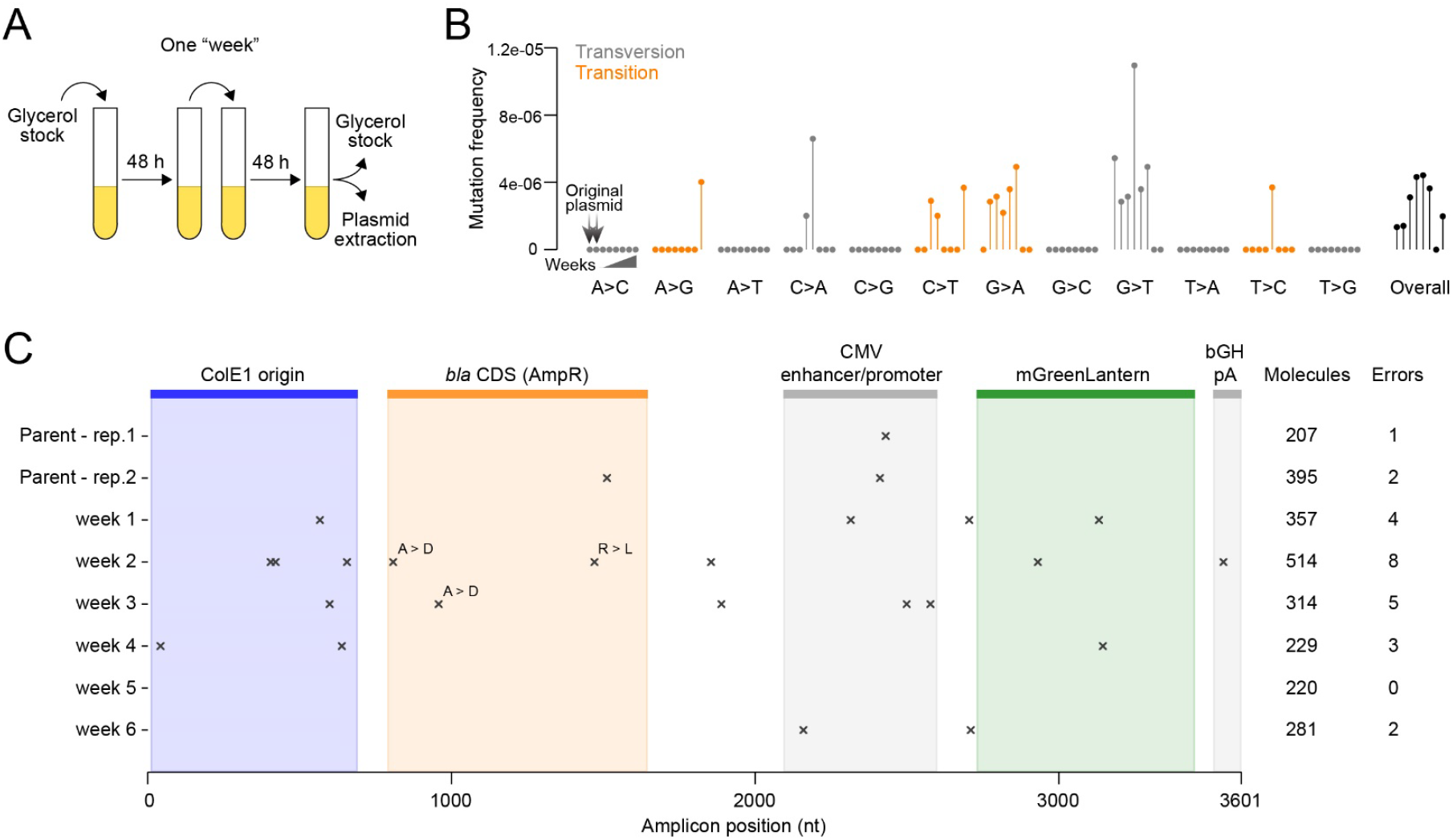
Quantifying plasmid mutations during bacterial culture. **A**) Schematic of plasmid culture protocol for a single “week”. The subsequent week begins with inoculation of LB broth using the glycerol stock established the prior week. Isolated plasmid DNA samples from each week, along with the original input plasmid, were UMI-labeled and sequenced. **B**) Rates of specific mismatch mutations observed in plasmids during long-term bacterial cell culture. Transition and transversion mutation types are distinguished by color. **C**) Position of detected mutations (mismatches and indels) in the longitudinal plasmid amplicons (left panel). The number of clusters analyzed and total errors relative to the template sequence are tallied (right panel).

### Quantifying PCR-induced mutations

Next, we sought to determine if PCR is responsible for introducing errors at a rate that would explain the prevalence of consensus sequences with errors we observed in the pooled plasmid experiment. A specific error introduced during PCR should not influence our consensus sequence generation unless it was present in at least half of a cluster’s reads. Thus, in theory, the mutation would need to be introduced during the first handful of cycles in order to influence our pipeline. To estimate an expected rate of error incorporation by DNA polymerase, we performed iterative PCR amplifications of a plasmid template using two high-fidelity DNA polymerases: Platinum SuperFi II and PrimeSTAR MAX. The plasmid template we chose was included in our previous plasmid mix experiment. After four rounds of PCR, each with 35 amplification cycles, we UMI labeled amplicons and sequenced products via nanopore. For PrimeSTAR MAX, we observed a significant association between PCR cycles and the frequency of mutations (F(1,3)=167.12; P=.001) (Fig. 4A). Using Platinum SuperFi II, however, PCR cycles were not significantly associated with the frequency of errors we detected (F(1,3)=2.58; P=.207). Mutation types were similar across both polymerases (Fig. 4B). Our estimated mutation rate for PrimeSTAR MAX (4.44×10^−^7 per bp per cycle; 95% CI: 3.35×10^−7^-5.54×10^−7^) is in line with reported error rates of high-fidelity DNA polymerases.^43,44^ Using the upper limit of the 95% confidence interval of our estimated mutation rate, we calculated the expected number of mutations introduced by two cycles of PCR and compared this result to the errors we observed in our plasmid pool experiment (Table S4). Even with this conservatively high estimate of the PCR-induced error rate, we observed more errors than would be expected to arise from two cycles of PCR (74 versus 20).

**Figure 4.**
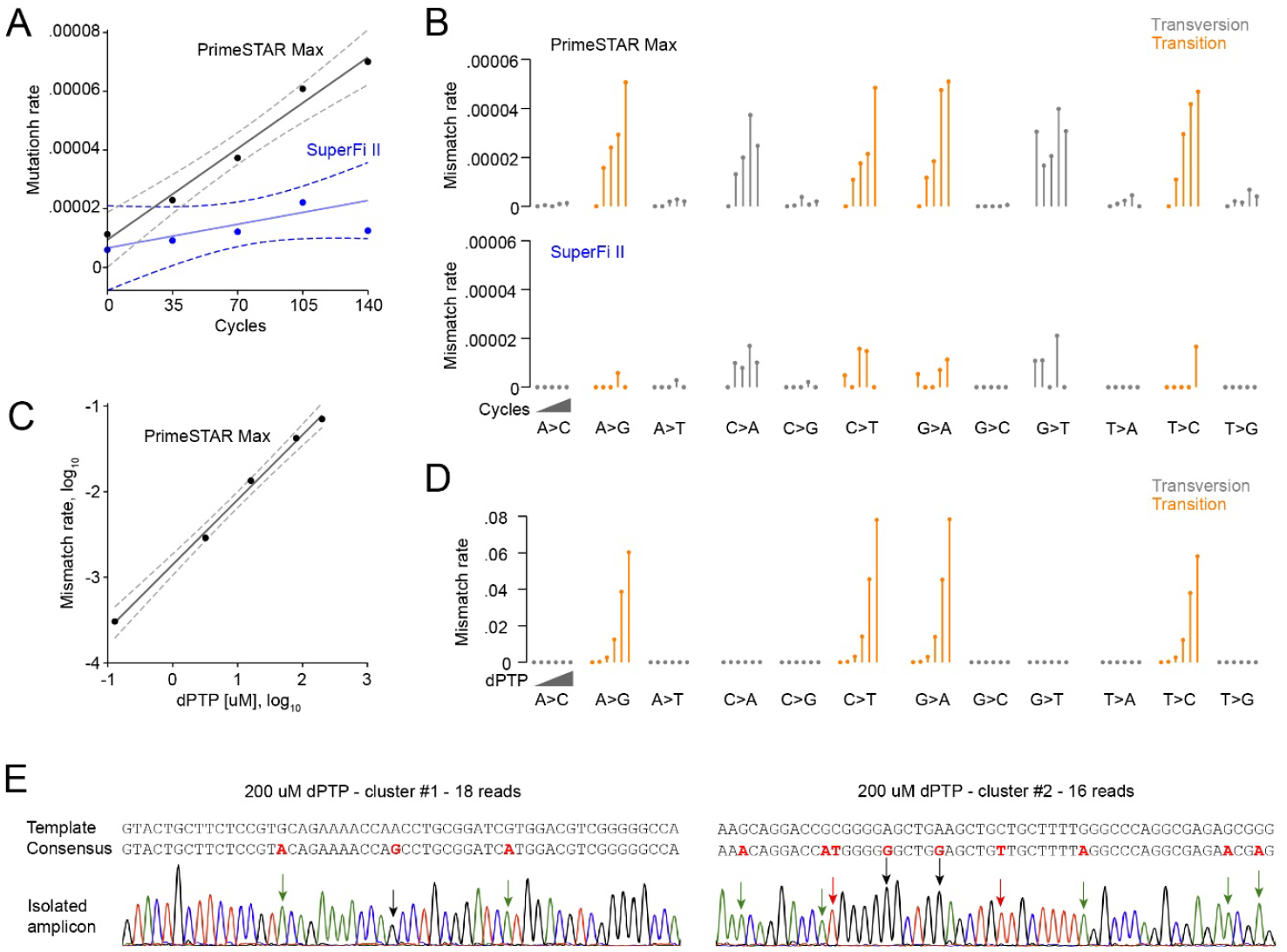
Quantifying the error rates of high-fidelity DNA polymerases. **A**) Scatterplot of the observed mismatch rate and the number of PCR cycles for two high-fidelity polymerase reagents. The solid and dashed lines represent the ordinary least squares regression fits and the associated 95% confidence intervals, respectively. **B**) Rates of specific mismatches observed across PCR iterations for PrimeSTAR (top panel) and SuperFi II (bottom panel) polymerase reagents. Transition and transversion mutation types are distinguished by color. **C**) Scatterplot of the observed mismatch rate as a function of dPTP concentration, both displayed on log_10_ scale. The solid and dashed lines represent the ordinary least squares regression fit and associated 95% confidence intervals, respectively. **D**) Rates of specific mismatches observed across increasing dPTP concentrations. Transition and transversion mutation types are distinguished by color. **E**) Transgene isolation from the 200 mM dPTP pool via PCR using UMIs as primer binding sites. Abridged cluster consensus sequences produced by the ConSeqUMI pairwise algorithm are shown below the template sequences. Chromatograms display the results of Sanger sequencing of the isolated molecules. Results for the full-length cluster #1 sequence are presented in Figure S6.

To rule out the possibility that we have underestimated the true PCR-induced error rate because of some upper limit of detection by our pipeline, we intentionally introduced random mismatch mutations in a plasmid fragment by performing PCR in the presence of increasing concentrations of the nucleoside triphosphate analog dPTP, which is readily utilized by *Taq* polymerase during DNA synthesis and triggers transition mutations after incorporation into newly synthesized strands.^45^ Resulting amplicons were then labeled with UMIs and further amplified via PCR prior to nanopore sequencing. Alignment of consensus sequences derived by ConSeqUMI to the original template sequence revealed a significant association between analog concentration and mutation frequency (F(1,3)=852.98; P=.0001) (Fig. 4C). The errors induced by dPTP were nearly exclusively transition mutations, in line with previous results (Fig. 4D).^45^ To assess the accuracy of consensus sequences derived by our pipeline, we again used UMI sequences as primer binding sites to isolate two individual molecules from the sample treated with the highest concentration of dPTP (200 µM). Sanger sequencing of the retrieved amplicons showed that ConSeqUMI derived the correct consensus for both (Fig. 4E, S6). These results suggest that UMI labeling of target molecules via degenerate primers followed by processing of nanopore data with ConSeqUMI can accurately detect mutational frequencies of at least 7% and that the observed frequencies of mutations in our plasmid pool experiment cannot be fully explained by PCR-induced errors during UMI labeling and were most probably present in the original plasmid preparations.

### Profiling recombinant adeno-associated virus genome variance

Having benchmarked our ConSeqUMI pipeline, we then designed three experiments to further showcase its utility: assessing genome integrity of recombinant adeno-associated virus (AAV) preparations, profiling genome repairs following CRISPR/Cas9-directed gDNA cleavage, and identifying severe acute respiratory syndrome coronavirus 2 (SARS-CoV-2) spike protein variants present in patient samples. Previous studies utilizing short-read sequencing technologies suggest that recombinant AAV genomes are produced and packaged at high fidelity, but recent reports have found that AAV preparations can contain large proportions of virions with partial genomes.^46–49^ To determine if ConSeqUMI can be used to monitor AAV preparation quality, we generated two recombinant AAV preparations using a published methodology.^36^ These recombinant genomes contained a U6-sgRNA expression cassette and an eGFP CDS under the control of either a CMV or EF-1α core promoter. UMIs were added to isolated genomes after an initial 30 cycles of PCR amplification with Platinum SuperFi II and amplicons were then further amplified prior to nanopore sequencing (Fig. 5A). Of 101 molecules sequenced from the CMV promoter preparation, 73 (72.7%) showed deletions, all of which were at least 1247 nt in length. Of the 73 genomes with deletions, 20 contained insertions ranging from 101-969 nt in length. Our earlier quantification of PCR-induced errors did not produce indels at this frequency or severity so these likely arise during recombinant virus production. Many of these inserted sequences could be mapped to plasmids utilized to produce the virions. Portions of two insertions mapped to separate introns in the human genome. Of the 28 full-length genomes, only 12 were the correct sequence (11.9% of mapped sequences), but the PCR amplification prior to UMI labeling could be responsible for some of the incorrect sequences. In contrast, the EF-1α core promoter preparation had 199 full-length genomes out of 232 sequenced (85.8%). The remaining 33 genomes had deletions of at least 1250 nt from the 1855 nt genome. Of these, 14 also contained insertions ranging from 32-581 nt in length. A total of 95 full-length genomes had the correct sequence (40.9% of mapped sequences).

**Figure 5.**
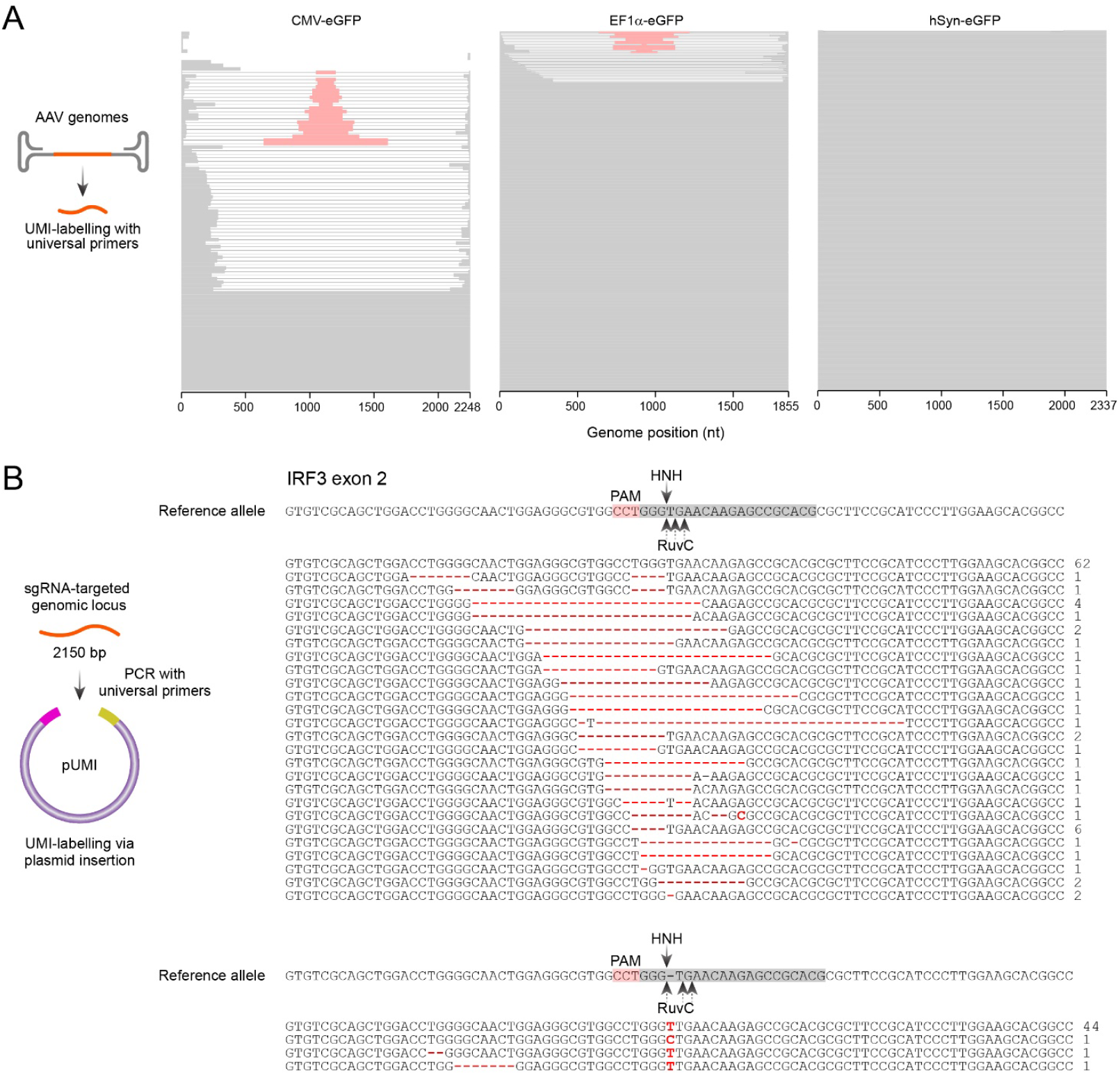
Evaluating recombinant AAV preparations and CRISPR/Cas9-induced indels. **A**) Recombinant AAV genome analysis. Stacked cluster sequences of CMV promoter (left panel), EF1a promoter (middle panel), and commercial (right panel) AAV preparations. Exogenous sequences inserted in truncated AAV genomes are shown in red. **B**) Using pUMI plasmid library to label amplicons derived from a sgRNA-targeted locus. The sequences represent each short deletion (top panel) and short insertion (bottom panel) at the sgRNA target region of human IRF3 exon 2 identified by the ConSeqUMI pipeline. Deleted and inserted nucleotides are shown in red. The reference allele is shown overlaid with the sgRNA target sequence (gray shadow), associated PAM sequence (red shadow), and the expected cleavage sites of the HNH and RuvC domains of SpCas9. The number of clusters with the abridged sequence are shown to the right of each sequence. The three observed insertions of longer size are shown in Figure S7.

After profiling our recombinant AAV samples, we next isolated genomes harboring a GFP expression cassette from an AAV preparation obtained from a commercial vendor (Addgene; pAAV-hSyn-eGFP) and prepared UMI-labeled amplicons for nanopore sequencing. With this commercial template, we were able to UMI-label genomes without prior PCR amplification. Of 292 clusters mapping to the template sequence, just two contained deletions (1945 nt and 1829 nt, respectively) from the center of the 2337 nt genome (Fig. 5A). Within the 290 full length genomes, we detected a total of 24 single-base mismatches (3.54×10^−5^ errors per nt sequenced) across 22 genomes, all of which were at unique locations. Thus, of 292 AAV genomes identified, 268 (91.8%) contained the desired sequence. The mismatch rate observed is eight-fold higher than we observed in the plasmid pool experiment (4.37×10^−6^ errors per nt sequenced), suggesting the majority of observed mismatches can be ascribed to viral genome replication machinery.

### Assessing Cas9-induced indels in human genomic DNA using plasmid-based UMI labeling

As an alternative to primer-based UMI addition, we developed a plasmid-based approach utilizing an acceptor plasmid with degenerate regions flanking a target insertion site (pUMI) (Fig. S7A). Following insertion of target molecules, resulting plasmids can be amplified by bacterial cells and an appropriate number of unique bacterial clones can be pooled and processed for nanopore sequencing. Advantages of this method compared to primer-based labeling are threefold: 1) finer control of the number of unique molecules to be sequenced, 2) prevention of both strands of the dsDNA receiving unique UMI pairs (see Fig. S1), and 3) preventing polymerase error introduction during UMI attachment. To test this approach, we utilized the pUMI library and ConSeqUMI to characterize insertions and deletions at a specific human genomic locus following cleavage by recombinant *Streptococcus pyogenes* Cas9 endonuclease (SpCas9) in HEK293T cells. We generated 2150 bp amplicons centered on the sgRNA target site at the human IRF3 locus and cloned these into the pUMI backbone (Fig. 5B). Of the 902 clusters with six or more reads, 164 mapped to the targeted locus. The majority of unmapped reads were derived from exon 4 of BAG3, the opposite strand of AMPD1, or intergenic regions on chromosomes 2, 5, and 7 (see deposited sequencing data). Within the mapped reads, 62 were unedited alleles and 102 contained deletions and/or insertions in proximity to the sgRNA target protospacer adjacent motif (PAM) (Fig. 5B, S7B). Of note, all detected insertions occurred precisely at the expected SpCas9 HNH domain cleavage site. Three alleles represented by six clusters contained large insertions of lengths 16, 58, and 190 bp, respectively (Fig. S7B). Five alleles represented by eight total clusters contained large deletions ranging in length from 149 to 668 bp. The most abundant repaired allele detected (44 clusters) contained an inserted thymidine at the HNH domain cleavage site, -3 relative to the PAM. This finding is in line with previous characterizations of non-homologous end joining following SpCas9-induced cleavage in mammalian cells in which the most prevalent repair was the insertion of a single nucleotide at the HNH domain cleavage site that matched the nucleotide of the -4 position relative the PAM.^50–52^ These results demonstrate the validity of our plasmid-based UMI labeling approach and that ConSeqUMI can be used to assess SpCas9-based genome editing in human cells.

### Isolating SARS-CoV-2 spike protein variants from human samples

The ability to identify virus genome variants is crucial for epidemiological monitoring and control of viral outbreaks. In the case of SARS-CoV-2, the sequence of the spike glycoprotein, which binds to host cell ACE2 receptors to facilitate viral entry, is used to classify virus strains. Single-molecule, long-read sequencing of the spike glycoprotein CDS with the ConSeqUMI platform could be used not only to identify virus strains present in patient samples, but also provide a snapshot of full-length spike protein heterogeneity that cannot be captured by short-read sequencing technologies. Using SARS-CoV-2 genomic RNA collected from four clinical samples, we first converted a 6162 nt region containing the full spike glycoprotein gene to cDNA and amplified the cDNA via PCR. Amplicons were then UMI-labeled, further amplified, then sequenced via Nanopore. For each cluster with 20 or more reads, we classified the viral lineage using Nextclade.^24^ For each patient sample, we identified a variety of spike glycoprotein sequences, suggesting intra-patient viral genomes are commonly heterologous, although we cannot rule out the unlikely possibility that all observed nucleotide variants were introduced by RT-PCR (see below). Three patient samples analyzed contained only A/J spike glycoprotein variants. However, in patient sample #1, we sequenced a total of 82 unique spike gene regions (622 total molecules, 616 were full length) and identified a mixture of delta variants: 73 variants of 21A/J (567 total molecules) and nine of 21I (55 total molecules). (Fig. 6A-C). All 55 sequenced spike proteins classified as 21I variants contained two nonsynonymous mutations - A222V and V367L - and a synonymous mutation at Y473, each absent from all 21A/J variants. A second synonymous mutation at I882, reverting to the Wuhan H-1 isolate, was seen in all 21I variants but in just three of 73 (4.1%) 21A/J variants. This sample also contained numerous spike glycoprotein variants containing the rare nonsynonymous substitution I834V that occurs predominantly in omicron lineage GJ.1.2.5 (97.3%).^53^ This substitution was present in 16 variants represented by 109 total clusters (17.5% of all spike genes sequenced in this sample). All spike genes sequenced from the other three samples contained only I834, encoded by the ATC codon. The existence of numerous recombinant SARS-CoV-2 lineages could indicate co-infection with separate lineages occurs in a proportion of infected individuals. We are aware of only a handful of previous reports of multi-strain SARS-CoV-2 infections, but since our experiment identified multiple strains present in one of four samples tested, this phenomenon may be more prevalent than presently appreciated.^54–58^ Alternatively, our sample with multiple strains may have resulted from cross-contamination during collection. Larger scale patient sampling will be required to assess the frequency of multi-strain infection in the human population.

**Figure 6.**
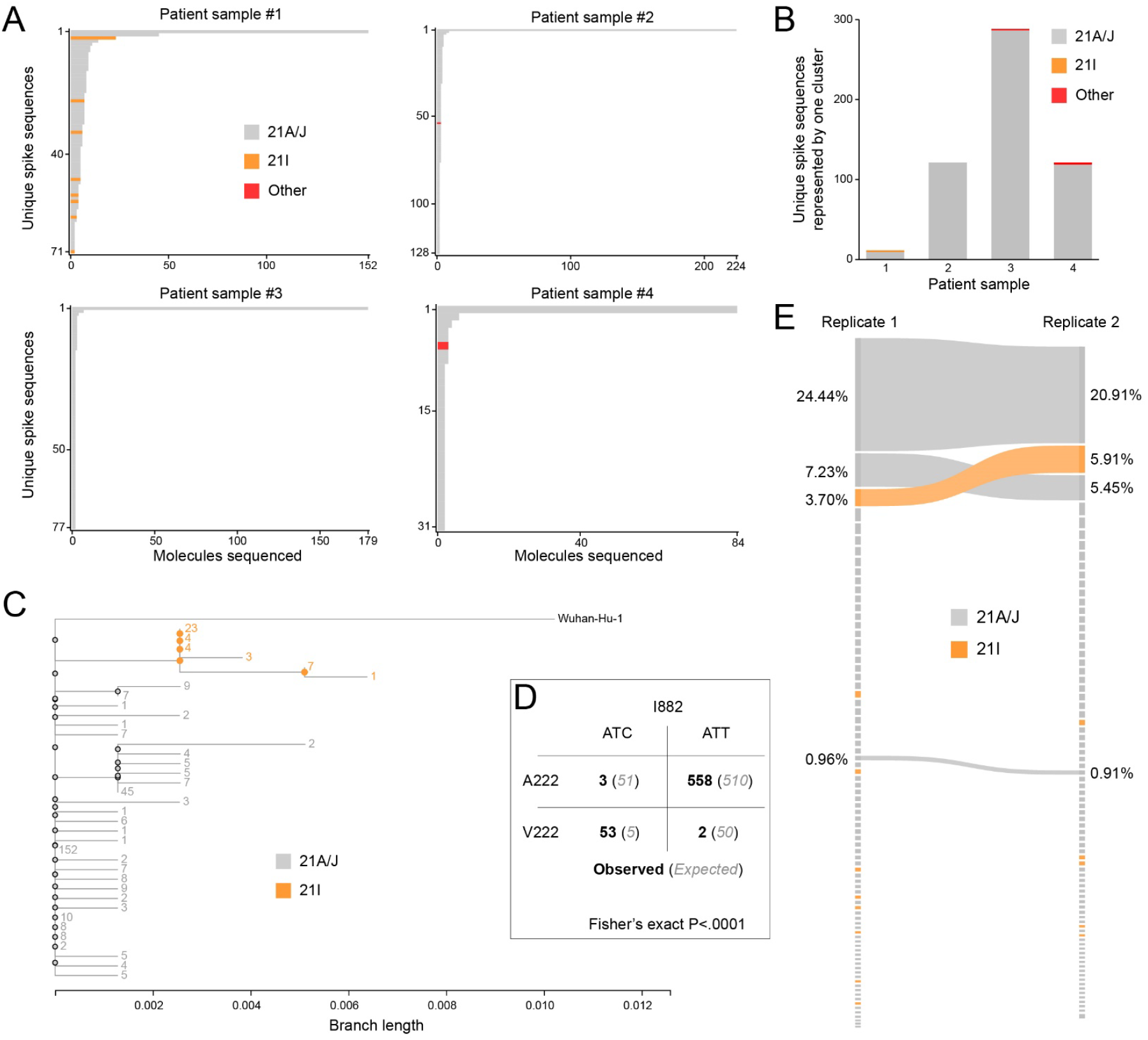
Profiling SARS-CoV-2 spike glycoprotein sequences in patient saliva. **A**) Unique SARS-CoV-2 spike protein CDS molecules represented by two or more clusters identified with ConSeqUMI in four patient samples. Strain identification was performed using Nextclade. **B**) Unique SARS-CoV-2 spike protein CDS molecules represented by exactly one cluster in the same four patient samples. **C**) Phylogram produced for translations of all 36 full-length spike protein CDS molecules lacking frameshift mutations in patient sample #1 and the original Wuhan isolate. The numbers adjacent to nodes and branches indicated the number of molecules sequenced. **D**) Contingency table of spike CDS molecules sequenced from patient sample #1 containing alanine or valine at position 222 and the ATC or ATT isoleucine codon at position 882. **E**) Sankey diagram of spike CDS sequences identified by repeated processing and sequencing of patient sample #1. Identical spike CDS molecules found in each replicate are connected by ribbons. The percentage of total molecules sequenced represented by each CDS is displayed by node height. Nodes and ribbons are colored according to strain classification assigned via Nextclade.

We detected a delta strain spike variant (represented by nine clusters) in sample #1 that did not contain the G142D mutation thought to be present in all delta variants. This particular lineage was classified as 21A/J by Nextclade, as it contained A222. Previous NGS sequencing results showing ‘wild type’ G142 in delta spike proteins were reported to be artifacts of the ARTIC V3 schema.^59^ Our results derived via Nanopore sequencing suggest that G142 does indeed exist in a proportion of delta spike glycoproteins and that G142D should not be used to define delta lineages.

One advantage of obtaining full length sequences from heterogeneous samples is that haplotype analyses can be performed across the entire sequenced region. For example, at amino acid position 882 of the spike glycoprotein from patient sample #1, 9.1% of genomes had a synonymous mutation (reverting to the Wuhan H-1 isolate ‘ATC’). Meanwhile, 8.9% of genomes contained the A222V nonsynonymous mutation, indicative of the 21I variant. The expected number of sequenced spike genes to contain both ATC at position 882 and a valine at position 222 was five, but we observed this combination in 53 of 616 full-length genes (Fisher’s exact P<.0001; Fig. 6D). This significant association between codons nearly 2000 nt apart could not have been identified using short-read sequencing approaches.

Because the identification of RNA viral genome sequences via Nanopore sequencing requires reverse transcription and because we found that amplification prior to UMI labeling was necessary for these particular samples, some mutations we detected may have been introduced during these procedures. We reasoned that performing independent RT-PCR and UMI labeling reactions on the same sample should produce the same spike CDS sequences if those sequences were true variants and not artifacts of the detection process. We therefore repeated the processing of patient sample #1 and sequenced the amplicons using a Flongle R10 flow cell. We sequenced a total of 220 molecules at a depth of 20 reads or more, compared to 622 molecules at the same depth in the original replicate. A total of four spike CDS sequences were identified in both replicates. However, these included the three most abundant sequences in each replicate and represented 33% and 36% of sequenced molecules (Fig. 6E). Because we are only subsampling input molecules at this sequencing depth, we speculate that many of the spike variants detected in just one of two replicates are in fact true variants. Nevertheless, our data indicates that the most abundant spike protein CDS variants detected with this method are true variants present in the input sample rather than artifacts produced by enzymatic conversion and amplification. Deeper, repeated sequencing analysis of patient samples with ConSeqUMI would reveal the depth of variant frequency in active infections.

## DISCUSSION

As long-read sequencing technologies continue to advance, the need for higher sequence accuracy remains a significant hurdle for their adoption across many applications. We have described the development and utilization of a novel computational platform for determining sequence identities for individual molecules present in heterogeneous mixtures of nucleic acids. ConSeqUMI software provides an all-in-one tool for the processing of UMI-labeled sequence reads collected and basecalled via the Oxford Nanopore sequencing platforms. Across multiple flow-cells and nanopore sequencing chemistries, we were able to obtain perfect single molecule sequence identity with as little as 20 reads per molecule. The nanopore platform is being continuously improved and greater sequencing accuracy of future chemistries and pores will further reduce the number of required reads per molecule to achieve similar results.

As we have shown here, determining the ground truth of a molecule’s sequence is impossible when PCR is required, as even high-fidelity commercial DNA polymerases will incorporate errors during amplification. Even with the low error rates we observed with two high-fidelity PCR systems, the origin of a given polymorphism detected by ConSeqUMI cannot be attributed to the original molecule with certainty. Nevertheless, we have demonstrated that ConSeqUMI does derive accurate consensus sequences for amplified molecules in heterogeneous pools. Furthermore, by replicating the amplification and UMI labeling processes on a single input sample, distinguishing true sample heterogeneity from potential technical errors becomes easier, as the probability of PCR amplification introducing identical error patterns during each replicate reaction becomes negligible.

Although UMI labeling and subsequent analysis with ConSeqUMI represents a significant improvement in the field of long-read sequencing, a number of technical limitations should be considered when implementing this approach. When performing UMI labeling, it is crucial to label an appropriate number of input molecules that will provide a suitable balance between sufficient sequencing depth and total input molecules sequenced at that depth. This process often requires troubleshooting, as labeling efficiency can vary depending on the stage of input dilution (before or after labeling) and the type of input sample. We recommend preparing libraries with the lowest molecule number that allows for gel extraction following amplification and to then inspect an aliquot of these libraries for read depth using a low throughput Flongle flow cell prior to running higher throughput flow cells. Ultimately, the number of molecules correctly identified will be limited by choice of sequencing platform and so care should be taken during experimental design. Regardless of platform, data collected will represent a subsampling of the input molecule population due to efficiencies of UMI labeling and the proportion of labeled molecules receiving adequate read depth. As such, molecule sequences with low occurrence in the data set are likely to have been more abundant in the input sample than other molecules that go undetected with this approach.

## DATA AVAILABILITY

Raw sequencing data and resulting consensus sequences produced by ConSeqUMI are deposited at the NCBI Gene Expression Omnibus under accession GSE288938. The ConSeqUMI package is available on GitHub (https://github.com/JGEnglishLab).

## ACKNOWLEDGEMENTS

The authors thank Dr. David O’Connor at the University of Wisconsin-Madison for providing SARS-CoV-2 RNA collected from infected individuals.^37^ The authors also thank Brett Milash at the Center for High Performance Computing at the University of Utah for critical insights.

## AUTHOR CONTRIBUTIONS

Conceptualization: A.M.Z. and J.G.E.; Formal analysis: A.M.Z. and C.W.C.; Funding acquisition: J.G.E.; Investigation: A.M.Z., A.N.G., K.E.R. and B.S.; Methodology: A.M.Z., C.W.C., B.S., and J.G.E.; Project administration: J.G.E.; Resources: J.G.E.; Software: C.W.C. and S.R.H.; Supervision: J.G.E.; Validation: A.M.Z.; Visualization: A.M.Z.; Writing – original draft: A.M.Z.; Writing – review and editing: A.M.Z. and J.G.E.

