## Supplemental Figures for "ConSeqUMI, an error-free nanopore sequencing pipeline to identify and extract individual nucleic acid molecules from heterogeneous samples"

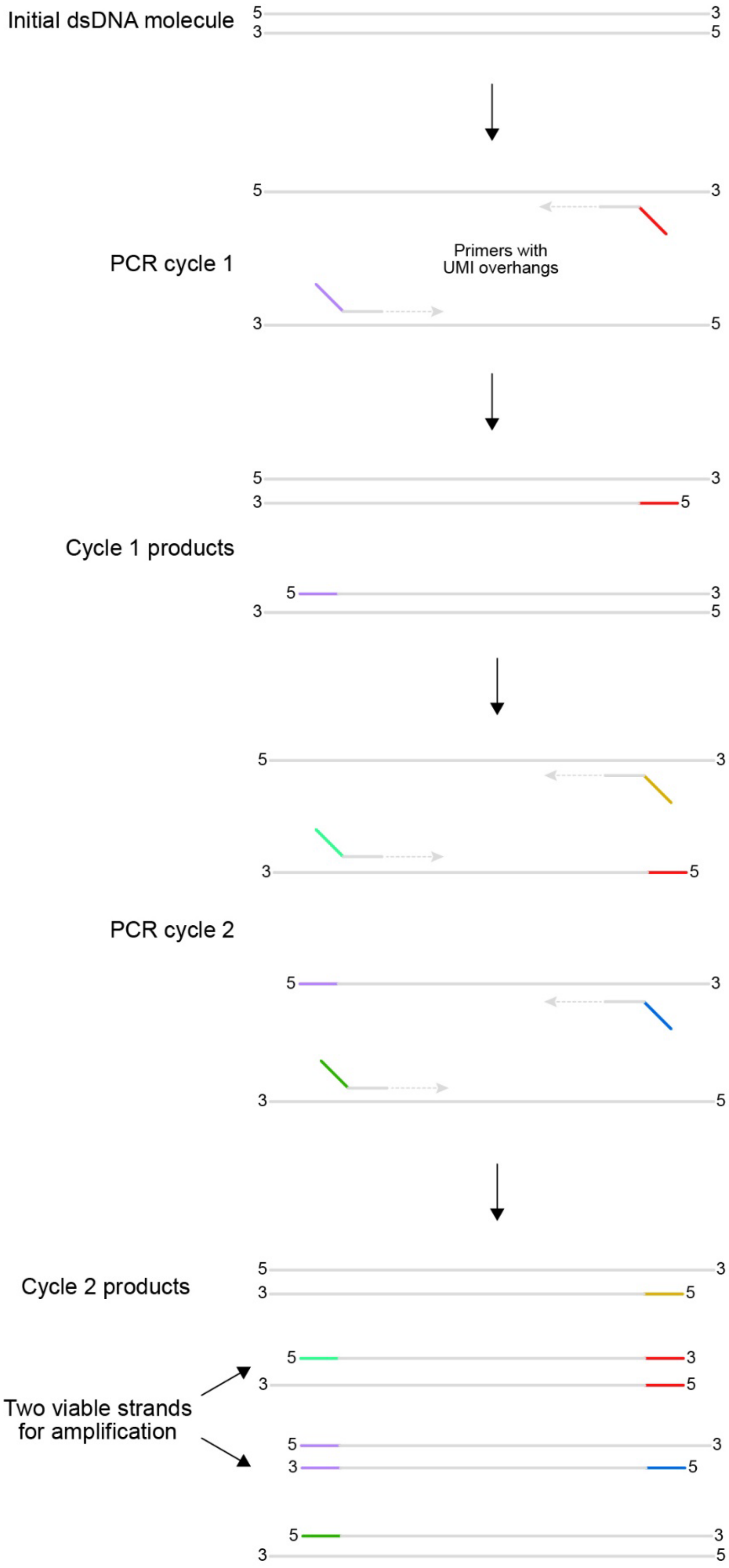


**Figure S1**. UMI labeling with PCR produces two daughter UMI pairs
Each input dsDNA molecule will produce two ssDNA strands that are able to be amplified by subsequent PCR using primers external to the UMIs. Each ssDNA has unique 5’ and 3’ UMIs.

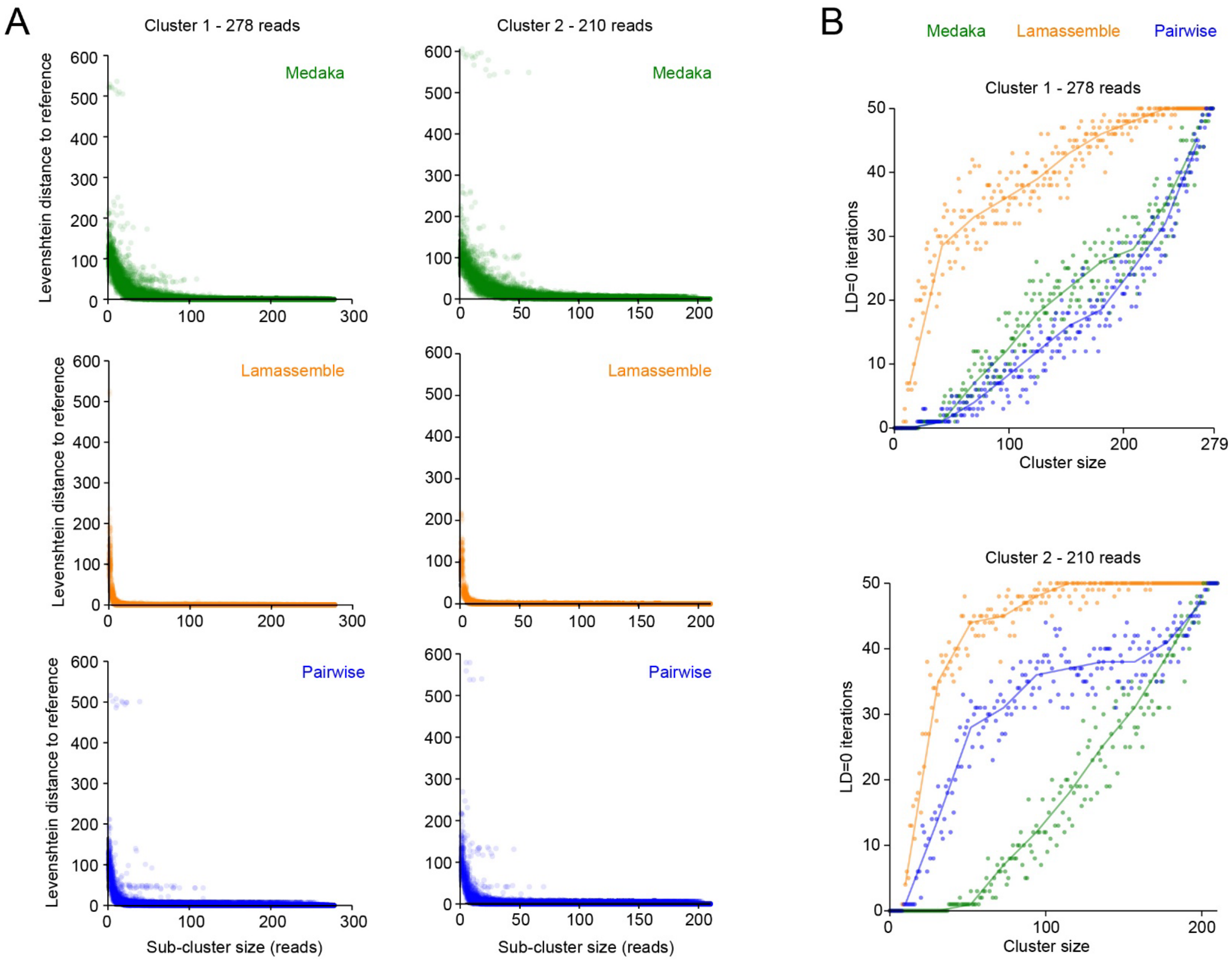


**Figure S2**. Benchmarking (barcode)

**A**) Benchmarking of each consensus generation algorithm available with ConSeqUMI for two clusters of the pcDNA3.1 pool experiment. For each sub-cluster size, the consensus generation procedures were repeated 50 times using randomly selected reads for each. Sub-cluster consensus sequences were then compared to the consensus generated by all available reads using Levenshtein distance. **B**) The number of iterations with a Levenshtein distance (LD) of zero at each cluster size used for consensus sequence benchmarking of two independent clusters. Lines represent the cross medians with band sizes of ten.

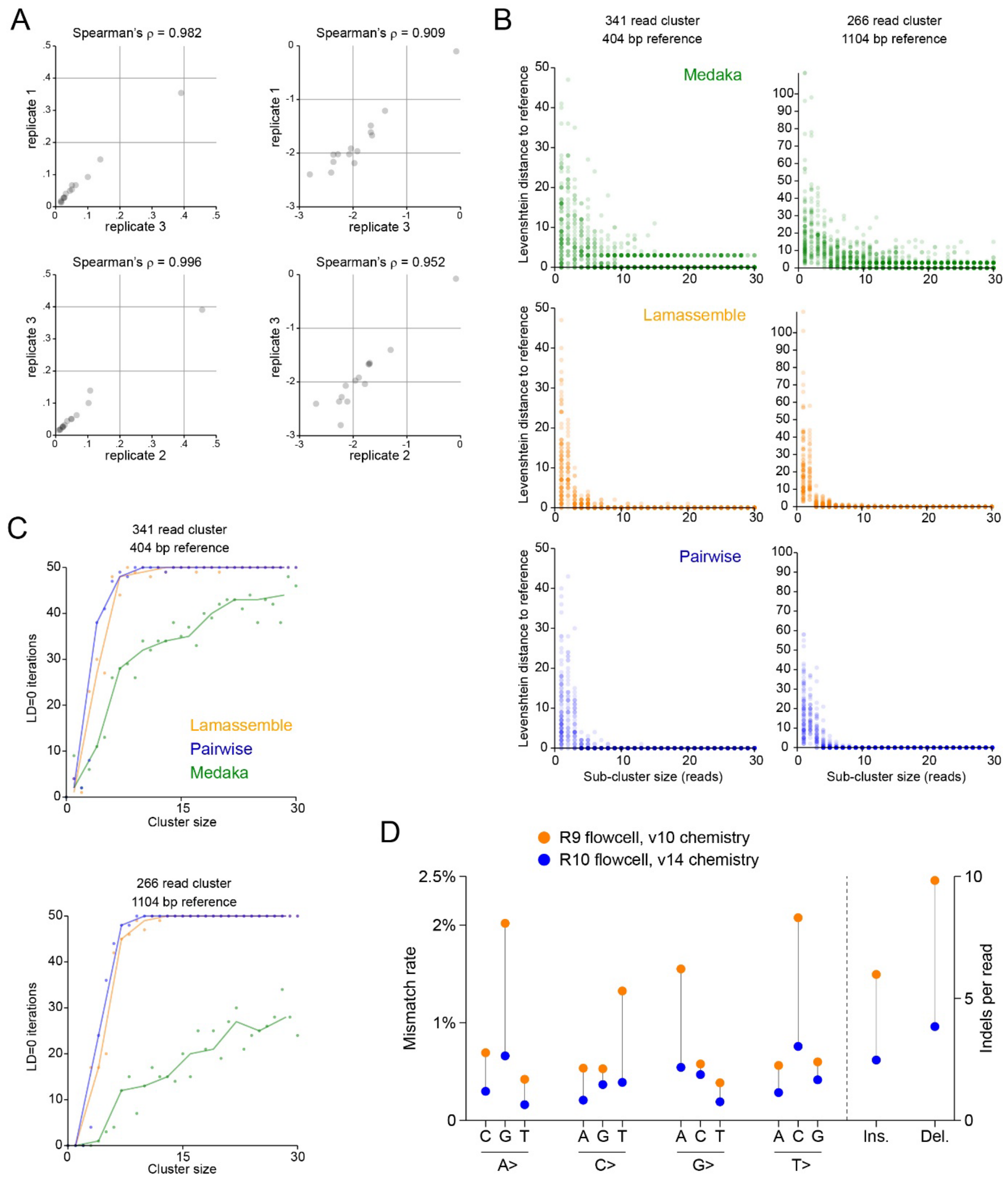


**Figure S3.** Benchmarking (pcDNA mix)
**A**) Replicate UMI-labeled plasmid pool samples were sequenced and clustered using ConSeqUMI. Primer-based UMI addition showed consistent cluster identity proportions across each of three replicates per input amount. **B**) Benchmarking of each consensus generation algorithm available with ConSeqUMI for two clusters of the pcDNA3.1 pool experiment. For each sub-cluster size, the consensus generation procedures were repeated 50 times using randomly selected reads for each. Sub-cluster consensus sequences were then compared to the consensus generated by all available reads using Levenshtein distance. **C**) The number of iterations with a Levenshtein distance of zero at each cluster size used for consensus sequence benchmarking of two independent clusters. Lines represent the cross medians with band sizes of ten. **D**) Comparison of nucleotide mismatch and indel rates between the R9 and R10 flow cells and associated reagent chemistries.

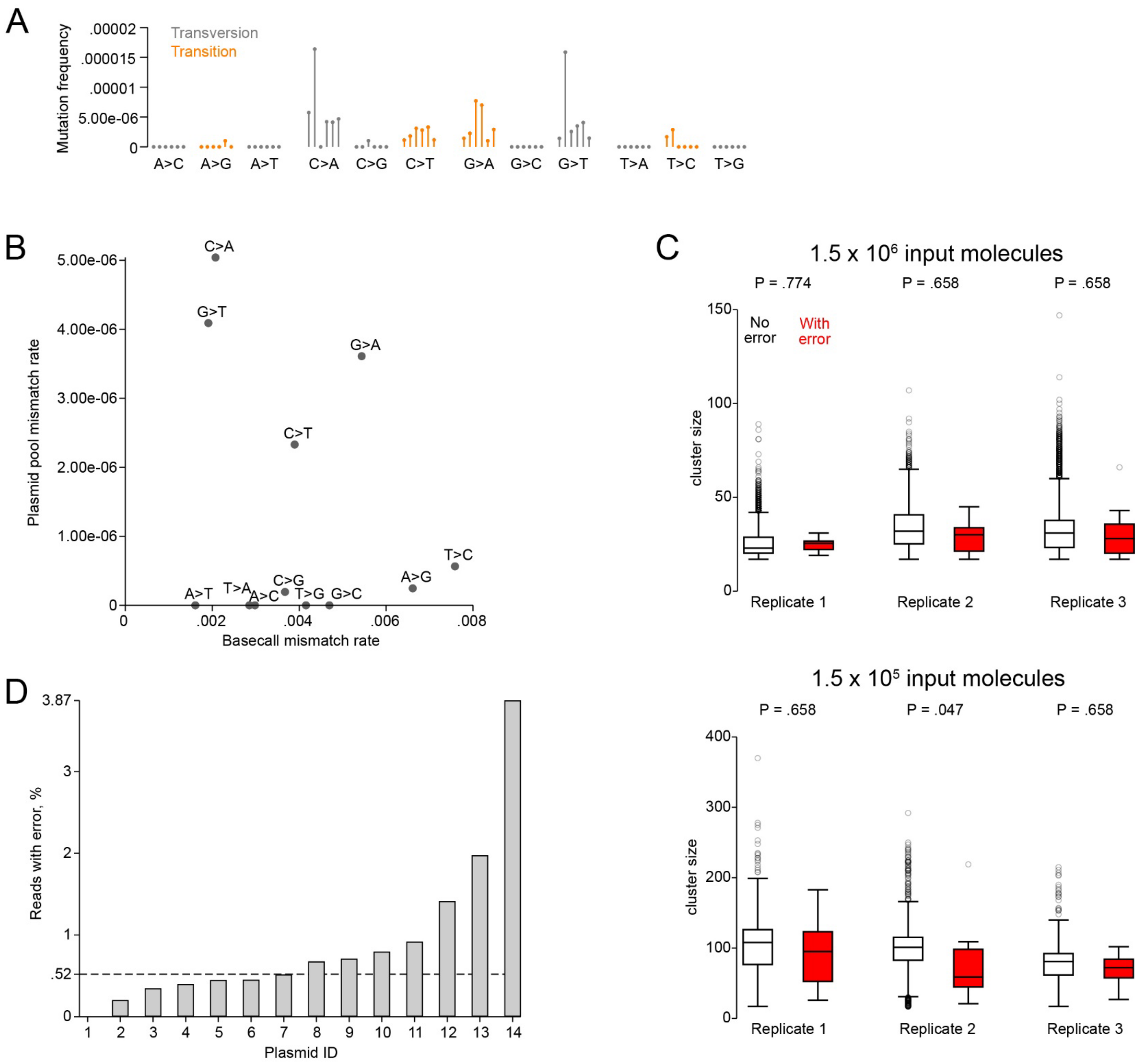


**Figure S4.** Plasmid sequence errors
**A**) Rates of specific mismatches observed for each input replicate. Transition and transversion mutation types are distinguished by color. **B**) Scatterplot comparing the mismatch error rates observed in the plasmid pool clusters and the baseball errors observed in individual sequencing reads. **C**) Comparison of cluster sizes for consensus sequences with (red) and without (white) errors (mismatch or indel) relative to the reference sequence. P-values represent Wilcoxon rank sum tests with multiple comparisons correction performed using the Sidak-Holm method.[1] Box lines represent 25th, 50th, and 75th percentiles. Whiskers represent upper and lower adjacent values as defined by Tukey.[2] Circles are outside values. **D**) The percentage of reads with at least one error (mismatch or indel) for each of the 14 plasmids in the pool. The dashed line indicates the overall percentage of reads with error (0.52%).

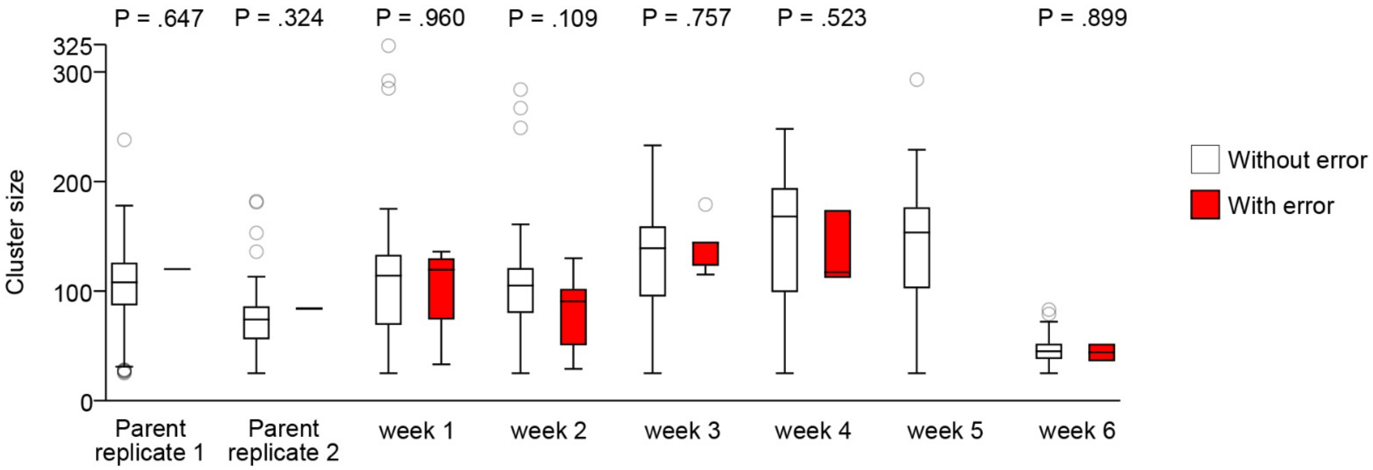


**Figure S5.** Longitudinal plasmid errors
Comparison of cluster sizes for consensus sequences with (red) and without (white) errors relative to the reference sequence. P-values represent Wilcoxon rank sum tests. Box lines represent 25th, 50th, and 75th percentiles. Whiskers represent upper and lower adjacent values as defined by Tukey.[2] Circles are outside values.

### 200 uM dPTP - cluster #1 - 18 reads

```
           1                                                                                                  100
 Template  AAGCAGAGCTCTCTGGCTAACTAGAGAACCCACTGCTTACTGGCTTATCGAAATTAATACGACTCACTATAGGGAGACCCAAGCTGGCTAGCGTTTAAAC
Consensus  AAGCAGAGCTCTCTGGCTAACTAGAGAACCCACTGCTTACTGGCCTATCAAAATTAATACAACTCACTATAGGGAGACCCAAGCTGGCTAGCGTTTAAAC
   Sanger                        AGAGAACCCACTGCTTACTGGCCTATCAAAATTAATACAACTCACTATAGGGAGACCCAAGCTGGCTAGCGTTTAAAC

           101                                                                                                200
 Template  TTAAGCTTGGTACCACCATGGCCCGCTCGCTGACCTGGCGCTGCTGCCCCTGGTGCCTGACGGAGGATGAGAAGGCCGCCGCCCGGGTGGACCAGGAGAT
Consensus  TTAAGCTTGGTACCACCATGACCCGCTCGCTGACCTGGCGCTGCTGCCCCTGGTGTCTGACGGAGGATGAGAGGGCCGCCGCCCGGGTGGACCAGAAGAT
   Sanger  TTAAGCTTGGTACCACCATGACCCGCTCGCTGACCTGGCGCTGCTGCCCCTGGTGTCTGACGGAGGATGAGAGGGCCGCCGCCCGGGTGGACCAGAAGAT

           201                                                                                                300
 Template  CAACAGGATCCTCTTGGAGCAGAAGAAGCAGGACCGCGGGGAGCTGAAGCTGCTGCTTTTGGGCCCAGGCGAGAGCGGGAAGAGCACCTTCATCAAGCAG
Consensus  CAACAGGATCCTCTTGGAGCAGAAGAAGCAGGACCGCGGGAAGCTGAAGCTACTGCTTTTGGGCCCAGGCGAGAGCGGGAAGAACACCTTCATCAAGCAG
   Sanger  CAACAGGATCCTCTTGGAGCAGAAGAAGCAGGACCGCGGGAAGCTGAAGCTACTGCTTTTGGGCCCAGGCGAGAGCGGGAAGAACACCTTCATCAAGCAG

           301                                                                                                400
 Template  ATGCGGATCATCCACGGCGCCGGCTACTCGGAGGAGGAGCGCAAGGGCTTCCGGCCCCTGGTCTACCAGAACATCTTCGTGTCCATGCGGGCCATGATCG
Consensus  ATGCGGATCATCCACGGCGCCGGCTACTCGAAGGAGGAGTGCAAGGGCTTCCGGCCCCCGGTCTACCAGAACATCTTCGTGTCCATGCAGGTCATGATCA
   Sanger  ATGCGGATCATCCACGGCGCCGGCTACTCGAAGGAGGAGTGCAAGGGCTTCCGGCCCCCGGTCTACCAGAACATCTTCGTGTCCATGCAGGTCATGATCA

           401                                                                                                500
 Template  AGGCCATGGAGCGGCTGCAGATTCCATTCAGCAGGCCCGAGAGCAAGCACCACGCTAGCCTGGTCATGAGCCAGGACCCCTATAAAGTGACCACGTTTGA
Consensus  AGGCCATGGAGCGGCTACAGATTCCATTCAGCAGGCCCGAGAACAAGCACCACGCTAGCCTGGTCATGAACCGGGACCCCTATAAAGTGACCACGTTTAA
   Sanger  AGGCCATGGAGCGGCTACAGATTCCATTCAGCAGGCCCGAGAACAAGCACCACGCTAGCCTGGTCATGAACCGGGACCCCTATAAAGTGACCACGTTTAA

           501                                                                                                600
 Template  GAAGCGCTACGCTGCGGCCATGCAGTGGCTGTGGAGGGATGCCGGCATCCGGGCCTGCTATGAGCGTCGGCGGGAATTCCACCTGCTCGATTCAGCCGTG
Consensus  GAAGCGCTACGCTGCGGCCATGCAGTGGCTGTGGAGGGATGCCGGCATCCGGGCCCATTATGAGCGTCGGCGGGAATTCCACCTGTTCGATTCAGCCGTG
   Sanger  GAAGCGCTACGCTGCGGCCATGCAGTGGCTGTGGAGGGATGCCGGCATCCGGGCCCATTATGAGCGTCGGCGGGAATTCCACCTGTTCGATTCAGCCGTG

           601                                                                                                700
 Template  TACTACCTGTCCCACCTGGAGCGCATCACCGAGGAGGGCTACGTCCCCACAGCTCAGGACGTGCTCCGCAGCCGCATGCCCACCACTGGCATCAACGAGT
Consensus  TACTACCTGCCCCACCTGGAACACATCACCGAGGAGGGCTACGTCCCCACAGTTCAGGACGTGCTCTGCAGCCGCATGCCCATCACTGGCATCAACGAGT
   Sanger  TACTACCTGCCCCACCTGGAACACATCACCGAGGAGGGCTACGTCCCCACAGTTCAGGACGTGCTCTGCAGCCGCATGCCCATCACTGGCATCAACGAGT

           701                                                                                                800
 Template  ACTGCTTCTCCGTGCAGAAAACCAACCTGCGGATCGTGGACGTCGGGGGCCAGAAGTCAGAGCGTAAGAAATGGATCCATTGTTTCGAGAACGTGATCGC
Consensus  ACTGCTTCTCCGTACAGAAAACCAGCCTGCGGATCATGGACGTCGGGGGCCAGAAGTCAGAGCGTAAGAAATGGATCCATTGTTTCGAGAACGTGATCGC
   Sanger  ACTGCTTCTCCGTACAGAAAACCAGCCTGCGGATCATGGACGTCGGGGGCCAGAAGTCAGAGCGTAAGAAATGGATCCATTGTTTCGAGAACGTGATCGC

           801                                                                                                900
 Template  CCTCATCTACCTGGCCTCACTGAGTGAATACGACCAGTGCCTGGAGGAGAACAACCAGGAGAACCGCATGAAGGAGAGCCTCGCATTGTTTGGGACTATC
Consensus  CCTCATCTACCTGGCCTCACTGAGTGAATACGATCAGTGCCTGGAGGAGAACAACCAGGAGAACCGCATGAAGGAGAGCCTCGCATTGTTTGGGACTATC
   Sanger  CCTCATCTACCTGGCCTCACTGAGTGAATACGATCAGTGCCTGGAGGAGAACAACCAGGAGAACCGCATGAAGGAGAGCCTCGCATTGTTTGGGACTATC

           901                                                                                               1000
 Template  CTGGAACTACCCTGGTTCAAAAGCACATCCGTCATCCTCTTTCTCAACAAAACCGACATCCTGGAGGAGAAAATCCCCACCTCCCACCTGGCTACCTATT
Consensus  CTGGAACTACCCTGGTTCAAAAACACATCCGTCATCCTCTTTCTCAACAAAACCGACATCCTGGAGGAGAAAATCCCCACCTCCCACCTGGCTACCTATT
   Sanger  CTGGAACTACCCTGGTTCAAAAACACATCCGTCATCCTCTTTCTCAACAAAACCGACATCCTGGAGGAGAAAATCCCCACCTCCCACCTGGCTACCTATT

           1001                                                                                              1100
 Template  TCCCCAGTTTCCAGGGCCCTAAGCAGGATGCTGAGGCAGCCAAGAGGTTCATCCTGGACATGTACACGAGGATGTACACCGGGTGCGTGGACGGCCCCGA
Consensus  TCCCCAGTTTCCGGGGCCCTAAGCAGGATGCTGAGGCAGCCAAGAGGTTCATCCTGGACATGTACACGAGGATGTACACCGAGTACGTGGACGGCCCCGA
   Sanger  TCCCCAGTTTCCGGGGCCCTAAGCAGGATGCTGAGGCAGCCAAGAGGTTCATCCTGGACATGTACACGAGGATGTACACCGAGTACGTGGACGGCCCCGA

           1101                                                                                              1200
 Template  GGGCAGCAAGAAGGGCGCACGATCCCGACGCCTCTTCAGCCACTACACATGTGCCACAGACACACAGAACATCCGCAAGGTCTTCAAGGACGTGCGGGAC
Consensus  GAGCAGCAAGAAGGGCGCACAATCCCGACGTCTCTTCAGCTACTACACATGTGCCACAGACACACAGAACATCCGCAAGGTCTTCAAGGACGTGCGGGAC
   Sanger  GAGCAGCAAGAAGGGCGCACAATCCCGACGTCTCTTCAGCTACTACACATGTGCCACAGACACACAGAACATCCGCAAGGTCTTCAAGGACGTGCGGGAC

           1201                                                                                              1300
 Template  TCGGTGCTCGCCCGCTACCTGGACGAGATCAACCTGCTGTGACTCGAGTCTAGAGGGCCCGTTTAAACCCGCTGATCAGCCTCGACTGTGCCTTCTAGTT
Consensus  TCGGTGTTCGCCCGCTACCTGGACGAGATCAACCTGCTGTAACTCAAGTCTAGAGGGCCCGTTTAAACCCGCTGATCAGTCTCGACCGTGCTTTCTAGTT
   Sanger  TCGGTGTTCGCCCGCTACCTGGACGAGATCAACCTGCTGTAACTCAAGTCTAGAGGGCCCGTTTAAACCCGCTGATCAGTCTCGACCGTGCTTTCTAGTT

           1301                                                                    1379
 Template  GCCAGCCATCTGTTGTTTGCCCCTCCCCCGTGCCTTCCTTGACCCTGGAAGGTGCCACTCCCACTGTCCTTTCCTAATA
Consensus  ACCAGCCATCTGTTGTTTGCCCTTCCCCCGCACCTTCCTTGACCCTGGAGGGTGTCACTCCCACTGTCCTTTCCTAATA
   Sanger  ACCAGCCATCTGTTGTTTGCCCTTCCCCCGCACCTTCCTTGACCCTGGAGGGTGTCACTCCCAC
```

**Figure S6.** dPTP cluster consensus accuracy
The consensus sequence derived by the pairwise method for a cluster from the sample treated with 200 uM dPTP aligned to the template sequence. The cluster was extracted from the sequenced pool using PCR with the cluster's UMIs as primers and the resulting amplicon was sequenced via Sanger. The sequencing results confirmed the accuracy of the derived consensus sequence.

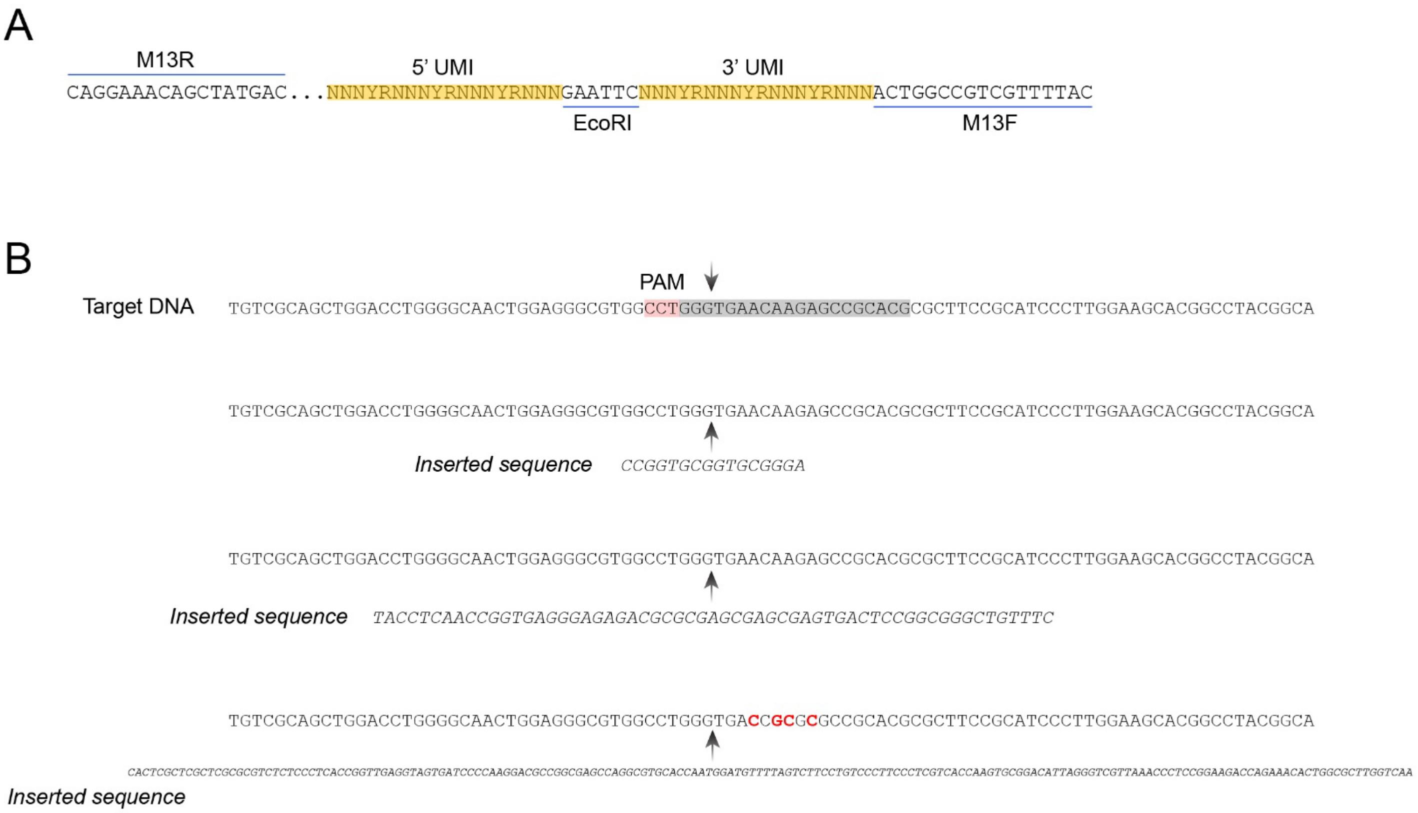


**Figure S7**. pUMI and IRF3 locus insertions
**A**) The pUMI plasmid library contains UMI regions flanking a single EcoRI recognition site where target molecules are inserted. Inserts with UMI labels can be liberated from either backbone for sequencing using HindII and SfoI restriction sites or by PCR using M13 primers. **B**) Three long insertions following SpCas9 cleavage of the human IRF3 locus identified by ConSeqUMI. All three insertions occurred at the predicted HNH domain cleavage site.
